# Spatial genome organization in nematodes with programmed DNA elimination

**DOI:** 10.1101/2025.10.23.684251

**Authors:** James R. Simmons, Tianchun Xue, Rachel Patton McCord, Jianbin Wang

## Abstract

Programmed DNA elimination (PDE) is a notable exception to genome integrity, characterized by significant DNA loss during development. In many nematodes, PDE is initiated by DNA double-strand breaks (DSBs), which lead to chromosome fragmentation and subsequent DNA loss. However, the mechanism of nematode programmed DNA breakage remains largely unclear. Interestingly, in the human and pig parasitic nematode *Ascaris*, no conserved motif or sequence structures are present at chromosomal breakage regions (CBRs), suggesting the recognition of CBRs may be sequence-independent. Using Hi-C, we revealed that *Ascaris* CBRs engage in three-dimensional (3D) interactions before PDE, indicating that physical contacts between break regions may contribute to the PDE process. The 3D interactions are established in both *Ascaris* male and female germlines, demonstrating inherent genome organization associated with the CBRs and to-be-eliminated sequences. In contrast, in the unichromosomal horse parasite *Parascaris univalens*, transient pairwise interactions between neighboring CBRs that will form the ends of future somatic chromosomes were observed only during PDE. Intriguingly, we found that *Ascaris* PDE, which converts 24 germline chromosomes into 36 somatic ones, induces specific compartmentalization changes. Remarkably, *Parascaris* PDE generates the same set of 36 somatic chromosomes, and the 3D compartment changes following PDE are consistent between the two species. Overall, our findings suggest that CBRs spatially demarcate the retained and eliminated DNA and may contribute to their spatial organization during *Ascaris* PDE. We also demonstrated that the 3D genome reorganization of the somatic chromosomes in these nematodes following PDE is evolutionary and developmentally conserved.

## Introduction

Genome integrity is crucial for faithful inheritance of genetic material. However, during the development of many species, programmed DNA elimination (PDE) removes a substantial portion of the genome, leaving daughter cells with reduced DNA content [1–6]. Although PDE is not present in most traditional model organisms (ciliates, for instance, do undergo PDE), it is being identified in a growing number of species from diverse phyla [7–14]. While PDE is gaining recognition as a significant biological process [15, 16], much remains unknown about its molecular mechanisms, particularly in metazoan species that exhibit PDE.

DNA double-strand breaks (DSBs) fragment chromosomes during PDE in ciliates, many nematodes, and some copepods [1, 2, 6]. During PDE, specific genomic locations are recognized, DSBs are generated, DNA break ends are healed by telomere addition or end joining, and specific regions of the genome are retained or eliminated. Although the molecular machineries for PDE appear divergent between the ciliates and nematode systems, both sequence-dependent and sequence-independent mechanisms are used for break site recognition [6, 7, 17, 18]. DSBs in the middle of the germline chromosomes, if healed with telomere addition, can lead to the partitioning of the chromosomes and drastic changes in the resulting somatic karyotypes [19]. However, the consequences of these karyotype changes on genome organization and gene expression are largely unexplored.

In the human and pig parasitic nematode *Ascaris* [20], we identified 72 chromosomal breakage regions (CBRs) at the boundaries of retained and eliminated DNA, with 48 terminal CBRs located near the chromosome ends – one per end across all 24 germline chromosomes [21]. Each CBR is in a 3-6 kb window where DSBs occur and new somatic telomere sequences are added to repair the broken ends [21, 22]. Sequence analysis revealed no significant motifs or sequence structures at these CBRs, suggesting a sequence-independent mechanism for their recognition [7]. Moreover, common histone marks, small RNAs, and R-loops are not associated with the CBRs [23, 24]. However, the chromatin at the CBRs becomes more accessible during PDE, indicating a specific mechanism may be involved [21, 22]. Similarly, the horse parasite *Parascaris* also possesses 72 CBRs where DSBs occur within a 3-6 kb window; these are internal CBRs confined in the euchromatic region of the single, large germline chromosome of 2.5 Gb of DNA [19].

The lack of sequence features associated with *Ascaris* and *Parascaris* CBRs prompted the search for sequence-independent mechanisms that might be involved in PDE. The eukaryotic genome is organized at a 3D level into folded structures at multiple length scales, involving both intrachromosomal (within the same chromosome) and interchromosomal (among different chromosomes) interactions [25, 26]. Changes in 3D genome organization have been implicated in numerous developmental processes, such as V(D)J and class switch recombination in lymphocytes [27, 28], zygotic genome activation during embryogenesis [29, 30], X-chromosome inactivation in mammals [31, 32], dosage compensation in *C. elegans* [33], antigenic variation and transcription in trypanosomes [34, 35], and neural development [36, 37]. We hypothesize that the 3D genome architecture may contribute to PDE by spatially organizing 1) the CBRs before and during PDE and 2) the resulting somatic chromosomes after PDE. Notably, in *Ascaris*, introducing random DSBs by treating with X-rays at PDE embryo stages did not lead to their repair through telomere addition [38]. This further suggests that the DSBs and telomere healing processes associated with *Ascaris* PDE are specific for PDE and may be confined within specific nuclear foci where the machineries for PDE could localize.

Recent advancements in chromosome conformation capture (Hi-C) have unveiled major principles of 3D genome organization, encompassing chromosome territories, compartments, topologically associating domains, point interactions, and acting through mechanisms such as phase separation and loop extrusion [25, 26, 39–43]. Here, to investigate the association between 3D genome organization and nematode PDE, we conducted Hi-C analyses on embryonic stages before, during, and after PDE. We found strong interactions between *Ascaris* CBRs prior to and during PDE that vanish after PDE, suggesting their potential involvement. Interestingly, newly formed somatic chromosomes undergo significant architectural changes following PDE, suggesting PDE leads to the establishment of new 3D genome organization in the somatic cells. In *Parascaris,* PDE partitions a single germline chromosome into 36 somatic chromosomes that share significant synteny with the 36 somatic chromosomes of *Ascaris*. The pattern of *Ascaris* somatic compartments is conserved in the horse parasite *Parascaris*, suggesting an evolutionarily preserved somatic 3D genome organization in ascarids. Together, our study provides insights into the dynamics of the 3D genome and nematode PDE and suggests a role for CBRs to spatially divide the retained and eliminated DNA and a potential function of PDE to influence the spatial reorganization of the somatic genome.

## Results

### Chromosomal breakage regions interact with each other during *Ascaris* DNA elimination

*Ascaris* early embryogenesis is slow (∼15 hours per cell cycle) and synchronous, and large quantities of embryos from discrete developmental staged can be collected [20], providing a unique opportunity to study the PDE process (Figure 1A) which occurs during the 4-16 cell embryonic stages [7]. To explore whether 3D genome organization plays a role in PDE, we performed Hi-C on *Ascaris* embryos at stages before (2-4 cells; 48 hr), during (4-8 cells; 60 hr), and after (32-64 cells; 5 days) PDE. The Hi-C data were mapped to an improved germline genome assembly that corrects previously mis-assembled regions (see Figure S1). The Hi-C heatmaps for individual chromosomes confirm the splitting of a single germline chromosome into multiple somatic chromosomes (Figure 1B). Across the genome, the Hi-C maps revealed a striking pattern of interactions prior to and during PDE among chromosome ends where the CBRs reside (Figure 1C, magenta arrows). These interactions are not due to mapping artifacts caused by repetitive sequences, since mapping the Hi-C data to a repeat-masked genome still produced the same pattern of interactions (Figure S2). The interactions are significantly higher than expected based on their genomic distance (Figure S3). Interactions also occur among CBRs located in the interior of chromosomes (Figure 1C, green arrows). Interestingly, the CBRs from the terminal and internal regions of the chromosomes form two spatially separated groups (Figure 1D-E). The two groups showed distinct interaction patterns, with terminal interactions forming either discrete point or streak interactions, while internal interactions appeared as long streaks (Figure 1F). Notably, these interactions, largely associated with the retained DNA, disappear after PDE, consistent with the timing of DNA elimination (Figure 1). Overall, the Hi-C data show that genomic regions associated with CBRs exhibit strong 3D interactions. These interactions are specifically associated with the CBRs, and they disappear immediately after PDE. Since the CBR is a unique feature associated with PDE, our data established a strong link between CBR interactions and PDE.

**Figure 1.**
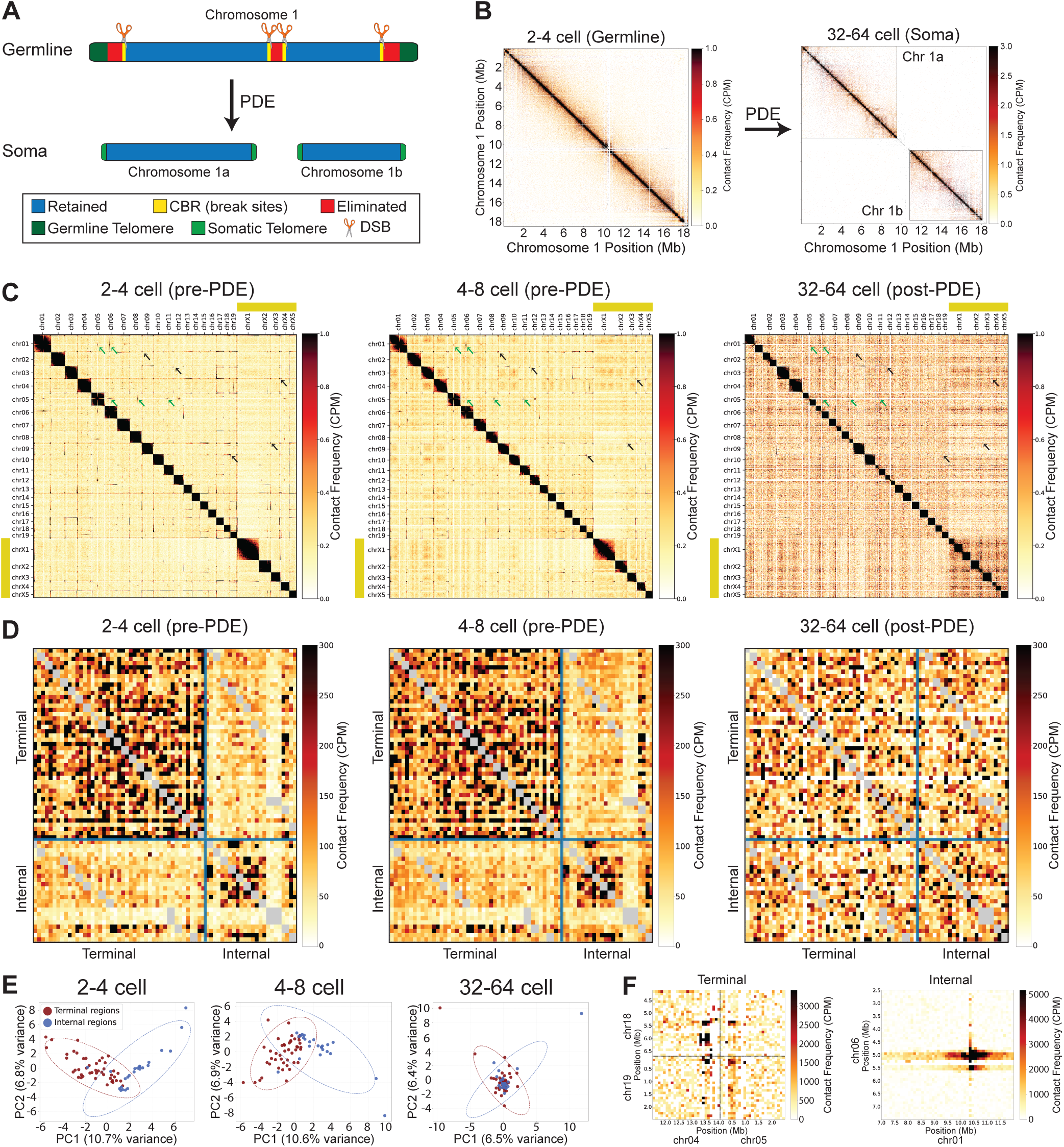
CBRs interact with each other during *Ascaris* PDE. **A.** PDE breaks one germline chromosome into two somatic chromosomes. Chromosome 1, one of the 11 *Ascaris* germline chromosomes that has DNA eliminated in the middle of the chromosome, is illustrated. **B.** Hi-C maps of chromosome 1 before and after PDE (2-4 cells and 32-64 cells, respectively) show the formation of the two somatic chromosomes following PDE. Contact maps were iteratively corrected (ICE method) and then normalized using counts per million (CPM). **C.** Hi-C heatmaps on *Ascaris* embryos from developmental stages before (2-4 cells, 48 hours), during (4-8 cells, 60 hours), and after (32-64 cells, 5 days) PDE are shown at 250 kb resolution. Before and during PDE, Hi-C interactions are observed between the ends of chromosomes where CBRs reside. Notably, these interactions are not observed after PDE at the 32-64-cell stage. Black arrows indicate examples of interactions at the ends of the chromosomes, while green arrows denote examples of interactions at the middle of the chromosomes. **D.** Hi-C heatmaps of 400-kb flanking each CBR reveal strong interactions among terminal (upper left blocks) and internal CBRs (lower right blocks) before and during the PDE. Note that these two clusters of CBR interactions are not visible in the post-PDE stage (32-64-cell embryos). **E.** Principal components analysis of CBR interactions reveals two distinct clusters before and during PDE and one cluster after PDE. **F.** Examples of terminal and internal CBR interactions binned at 100 kb resolution.

### CBR interactions are established during germline development

We next asked when the CBR interactions are formed. We performed Hi-C on male and female germline tissues and on 1-cell zygotes (0 hr), stages not associated with PDE. Surprisingly, we found that interactions among CBRs were already present in these germline tissues (Figure 2). Thus, these long-range CBR interactions are established well before PDE and may be an intrinsic feature associated with the CBRs. Interestingly, the patterns of 3D interactions at the ends of chromosomes exhibited variations among these germline stages (Figure 2). In the testis and 1-cell zygote, the ends of certain chromosomes are observed to interact with the entire length of many other chromosomes, resulting in stripes in the Hi-C maps (Figure 2A and 2C). In contrast, in the ovary, most interactions are more confined to the ends of the chromosomes, as indicated by the dots in the Hi-C maps (Figure 2B). Thus, while our data demonstrate an overall consistency in the CBR interactions throughout germline development, they also reveal stage-specific variations in the 3D genome interactions.

**Figure 2.**
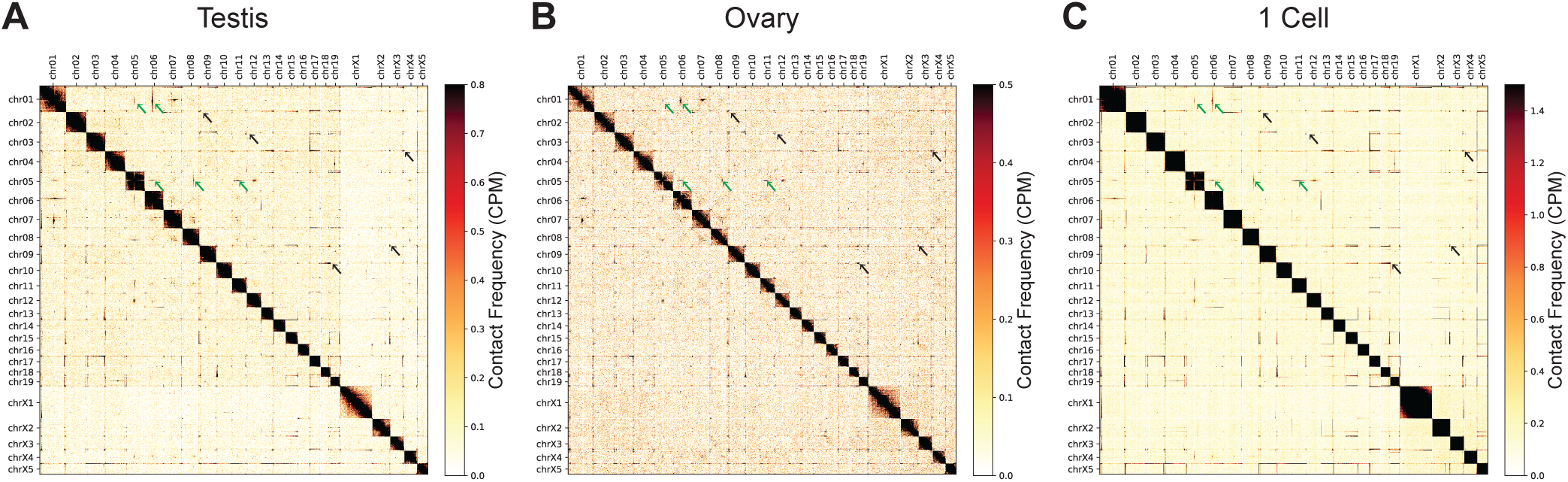
CBR interactions are established during germline development. Interactions between chromosome ends in *Ascaris* were present in the testis (**A**), ovary (**B**), and 1-cell zygote (**C**). Values were normalized using counts per million (CPM). Note the differences in the interactions as shown in dots (Ovary) vs. stripes (Testis and 1-cell zygote). Black and green arrows point to end-to-end and internal-to-internal interactions, respectively.

### CBRs reside at the boundaries of the spatially organized chromosome ends

To identify the precise spatial boundaries of the CBR interactions among chromosomes, we determined *trans*-interaction (interchromosomal) frequencies surrounding the CBRs. Due to the short DNA for many eliminated regions adjacent to CBRs (average 317 kb, median 197 kb), we chose to examine interchromosomal interactions of all terminal CBRs using 100 kb in the eliminated regions and 500 kb in the retained regions (Figure 3A). This analysis revealed a significant enrichment of *trans* interactions within ∼200 kb of retained DNA immediately adjacent to the CBRs near the ends of chromosomes (Figure 3A). Strikingly, CBRs coincide with the boundaries of the high and low interchromosomal interaction regions (Figure 3A, vertical line), suggesting these strong interactions may spatially demarcate the retained and eliminated sides of the DNA. Interestingly, this spatial separation is not detected for CBRs in the middle of the chromosomes (Figure S4), indicating that the stronger interchromosomal interactions for terminal CBRs may be linked their proximity to the chromosome ends.

**Figure 3.**
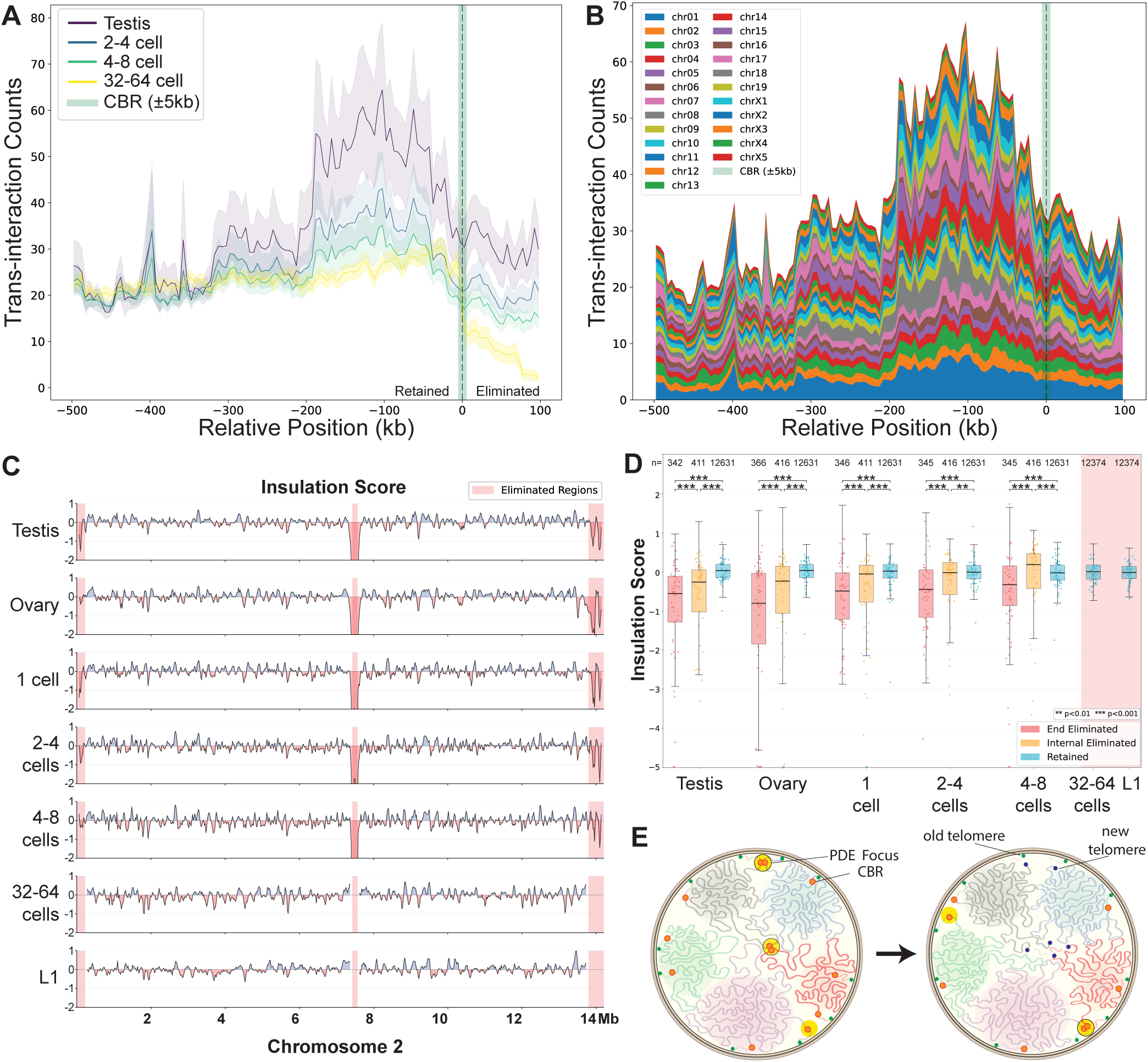
Interactions of CBR-adjacent regions are associated with PDE. **A.** CBR-adjacent regions have high interchromosomal interactions. The CBRs are highlighted in green with a vertical dashed line and designated as 0. The 500-kb retained side is to the left, and the 100-kb eliminated side is to the right. *Trans* interaction frequencies surrounding the CBRs at the ends of chromosomes are shown. Interactions among chromosomes were aggregated for each CBR-adjacent region using a 5-kb bin size. Meta-profiles for all analyzed CBRs are displayed as mean (solid lines) ± one standard error of the mean (SEM, shaded areas). **B.** All chromosomes contribute to the CBR interactions observed in Figure 3A. The aggregated CBR *trans*-interaction for the testis was broken down by contributions from each chromosome (ordered and color-coded). **C.** CBR regions are highly insulated in the germline and early embryos through PDE. Insulation score is shown for each developmental stage for chromosome 2. Less insulated regions (positive values) are shown in blue, more insulated regions (negative values) are shown in red. Eliminated regions are highlighted in red. c = cell(s). **D.** The end eliminated regions are highly insulated. The whole genome is broken down by retained (blue), end eliminated (red), and internal eliminated (orange) regions. Insulation score values using a 100-kb window size are plotted for each developmental stage in a sliding window size of 5 kb. Post-PDE stages on the right are highlighted in red (note the lack of eliminated DNA data in post-PDE). Data points below -5 are shown as hollow circles. Statistical significance was calculated using a two-sided Mann-Whitney U test (*** p<0.001 and ** p<0.01). **E.** Interaction of CBRs at PDE foci in the nucleus. A 3D genome interaction model based on Hi-C maps. Shown are the dynamic changes of the CBRs (red circles) in five chromosomes (five colored threads in their chromosome territories). PDE foci are shown in big yellow circles. Germline (old; green) and somatic (new; blue) telomeres are illustrated. Depicted are two CBRs interacting within a PDE focus (left, top), where a programmed DSB occurred. After DSB formation, new telomeres are added to the retained DNA, which then dissociates from the PDE foci (right, top). A newly formed CBR interaction is also shown (right, bottom). Meanwhile, internal CBRs interact among themselves apart from terminal CBRs, leading to drastic karyotype changes after elimination.

To further distinguish whether the elevated interactions in the CBR-flanking regions were driven by a few selected chromosomes or across all chromosomes, we analyzed the contribution of individual chromosomes. Each chromosome contributed considerably to the aggregated interactions (Figure 3B), suggesting a shared mechanism of 3D genome organization among these chromosomes. Interestingly, the interchromosomal interactions for CBR-adjacent retained regions are dynamic during embryo development, with the highest interactions observed in the early embryos; they gradually decrease throughout embryonic development and become minimal by L1 post-PDE (Figure 3A). Collectively, these results indicate that the chromatin architecture associated with CBRs employs long-range *trans* interactions prior to PDE. These interactions may play a key role in defining the boundaries of 3D genome domains that coincide with the CBRs, thus spatially separating the retained and eliminated DNA.

### Short-range interactions and 3D insulation associated with PDE

To explore whether short-range 3D interactions might also influence PDE, we calculated insulation scores across the genome. Insulation scores quantify the degree to which a given region interacts with its local genomic neighborhood. Dips in insulation score can therefore be interpreted as stronger local boundaries across which fewer interactions occur [33, 44]. We identified many genomic regions with dynamic insulation patterns during development (Figure 3C-D). Notably, out of the 44 completely assembled eliminated regions exceeding 100 kb that allow robust insulation analysis, 35 (80%) consistently exhibited boundary-like dips in insulation score at or near their respective CBRs before the PDE, while six (14%) demonstrated similar drops in insulation score in adjacent retained regions (Figure S5). Conversely, only one eliminated region (chrX5) did not show any sign of boundary-like insulation. Interestingly, we observed a general increase in insulation scores across eliminated regions up to their elimination at the 4-8-cell stage (Figure 3D). This coincided with the incorporation of the eliminated DNA fragments into micronuclei [21], which separate these fragments from the rest of the genome, thus may increase their local interactions. Overall, these findings suggest that increases in local boundary strength and 3D insulation are a common feature of *Ascaris* CBRs associated with PDE.

After PDE, the retained regions adjacent to CBRs become less insulated as indicated by the increasingly positive insulation scores, extending beyond 1 Mb (Figure 3C-D). The new chromosome ends also exhibit decreased gene expression, despite showing higher chromatin accessibility measured by ATAC-seq [22]. Consistently, the heterochromatic histone mark H3K9me3 was observed to accumulate at the new chromosome ends after PDE (Figure S6). Overall, this decoupling of accessibility and expression may reflect the repositioning of chromatin domains after PDE liberates the ends of their chromosomes and also splits some germline chromosomes.

Collectively, our data indicate that the CBRs interact with each other before and during *Ascaris* PDE and that CBRs are spatially delimiting the retained and eliminated DNA. The CBRs from the terminal and internal chromosomes form two spatially separated clusters. These 3D interactions of CBRs may play a role in the PDE process by bringing them to nuclear foci that may have accumulated molecular machinery essential for DSBs and telomere addition (PDE foci; see Figure 3E and discussion). Alternatively, the interactions of the CBRs themselves may act as a platform to recruit CBR- and/or PDE-associated proteins to the break sites.

### 3D chromosomal changes following programmed DSBs for PDE

In the previous sections, we have described the 3D interactions of CBRs and their potential association with the mechanisms of PDE. Below, we explore the 3D genome changes as a consequence of PDE. Although PDE coincides with the timing of embryogenesis, we will focus on 3D genome changes surrounding the DSBs that are specifically associated with PDE. Our previous analyses examined the insulation in retained genome regions adjacent to CBRs to determine if 1) the local 3D interactions had changed after PDE (Figure 3C) and 2) how the broad insulation patterns vary between the retained and eliminated regions leading up to PDE (Figure 3D). Here, to study the local insulation in response to the DSBs, we performed Hi-C analysis on several staged embryos post-PDE (Figure 4) and determined the insulation scores specifically for CBRs. Because the eliminated DNA is no longer available in the genomes of post-PDE stages, we only examined the retained region within 100 kb sequences around CBRs. These regions became more insulated from the 1-cell zygote to PDE stage (4-8 cells) but became less insulated after PDE (32-64 cells) (Figure 4A). This suggests that DSBs may liberate the spatial constraints adjacent to the CBRs during PDE, thus changing their local genomic contacts. Thus, our Hi-C data defined a consistent pattern of local 3D genome response to DSBs post-PDE.

**Figure 4.**
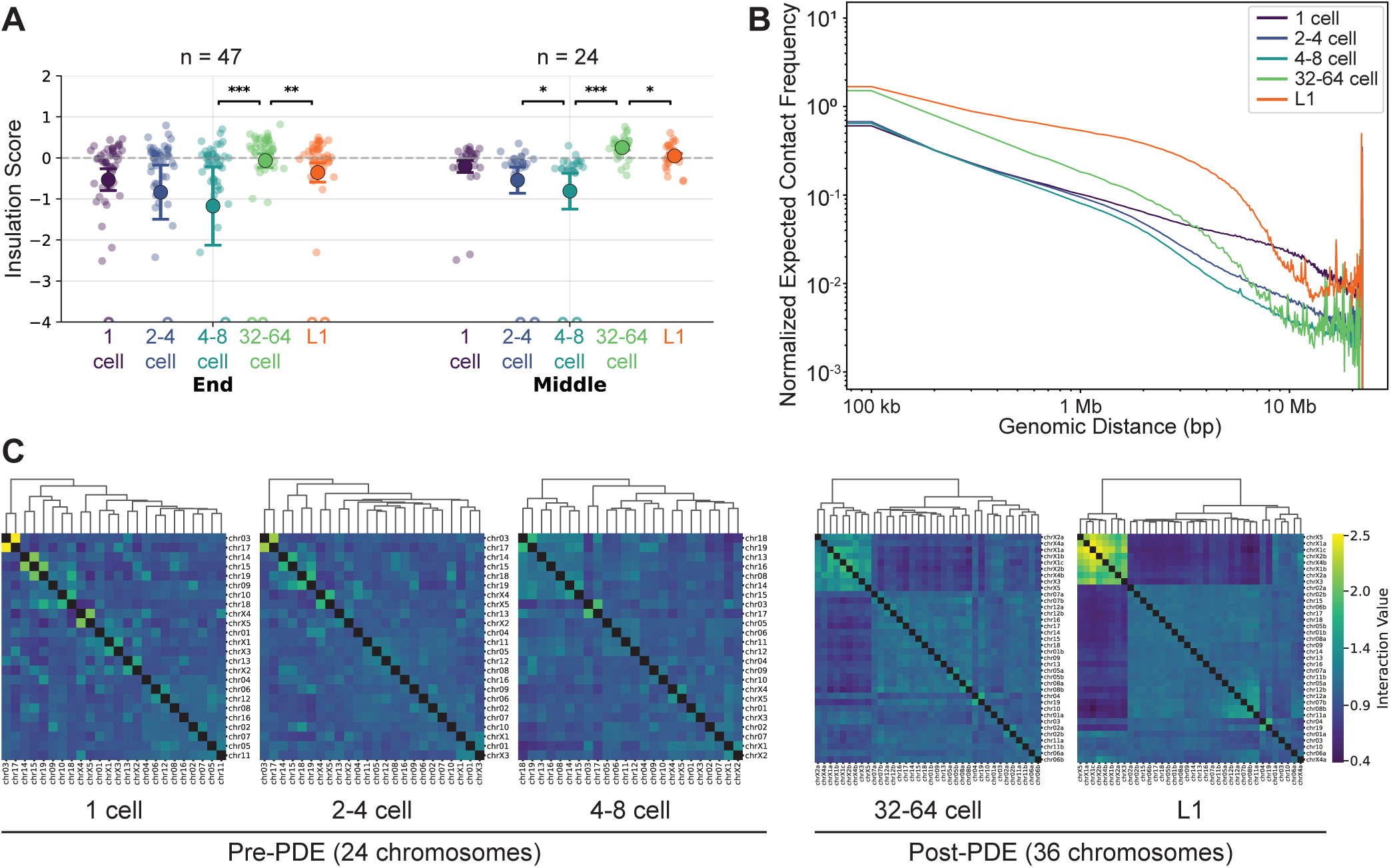
3D chromosomal changes during embryogenesis and in response to programmed DSBs. **A**. Dynamic changes of insulation for retained ends of chromosomes. Insulation scores for 100 kb regions of retained DNA proximal to the end (47 sites; left) and middle (24 sites; right) of the chromosomes were determined using Hi-C contact matrices at 20-kb resolution with a 100-kb insulation square size. Shown are mean values with bars representing one standard error. Each dot represents the end of a somatic chromosome (or future somatic chromosome in pre-PDE stages). Statistical significance was calculated using a two-sided Mann-Whitney U test (*** p<0.001, ** p<0.01, and * p<0.05). Data points lower than -4 are shown as empty circles at the lower boundary. **B.** Changes in middle and long-range Hi-C interaction. Distance-decay curves for whole-genome Hi-C interaction data at different developmental stages. Normalized expected contact frequency (y-axis) is plotted against genomic distance (x-axis) on a log-log scale. **C.** Changes in the 3D interaction among chromosomes. Chromosomal cluster analysis was performed at each developmental stage. Pre-PDE genomes are shown with 24 chromosomes, while post-PDE genomes are shown with 36 chromosomes. Strong clusters emerge after PDE as observed in the group of sex chromosomes (top-left corners) in the 32-64-cell embryos and L1 larvae.

### Long-range interaction changes and chromosome clustering through PDE

To assess the impact of PDE on intrachromosomal contacts, we analyzed distance decay curves across different stages (Figure 4B). In the 1-cell zygote, interaction frequency exhibits a power-law decay diminishing around 23 Mb, which aligns with the largest germline chromosome size (chrX1 = 23.1 Mb). Short-range interactions (<100 kb) increase after PDE, coinciding with the observed increase in local structures seen in the Hi-C heatmap (Figure 1). This suggests the establishment of short-range regulatory interactions throughout the genome following PDE. However, mid-range interactions (100 kb to 1 Mb) decline during early embryogenesis through the PDE stages but then rise again post-PDE within somatic chromosomes as embryogenesis proceeds, peaking in L1 larvae. Conversely, long-range interactions (>1 Mb) decrease after PDE, consistent with the overall reduced chromosome sizes (average size: germline = 12.8 Mb and soma = 7 Mb). These observations indicate that PDE contributes to the 3D architecture changes by altering both chromosome numbers, sizes, and the long-range interactions, facilitating the establishment of the 3D genome in the somatic nucleus.

To determine how chromosomes spatially reorganize following PDE, we clustered chromosomes at each developmental stage based on Hi-C contact frequencies. In the 1-cell zygote and 2-4-cell embryos, interactions among pre-PDE chromosomes are broadly uniform (Figure 4C). After PDE, the somatic chromosomes exhibit differential spatial organization. A key feature after PDE is the clustering of sex chromosomes: at 32-64 cells, the sex chromosomes form a distinct group separated from the autosomes. This segregation of the sex chromosomes becomes more pronounced in larvae L1. In contrast, for autosomes, they are organized into several large, distinct clusters, indicating progressive establishment of preferential interactions between chromosome territories (Figure 4C). These data suggest coordinated compartmentalization changes (see also Figure 5) may contribute to the formation of the somatic chromosome territories after PDE.

**Figure 5.**
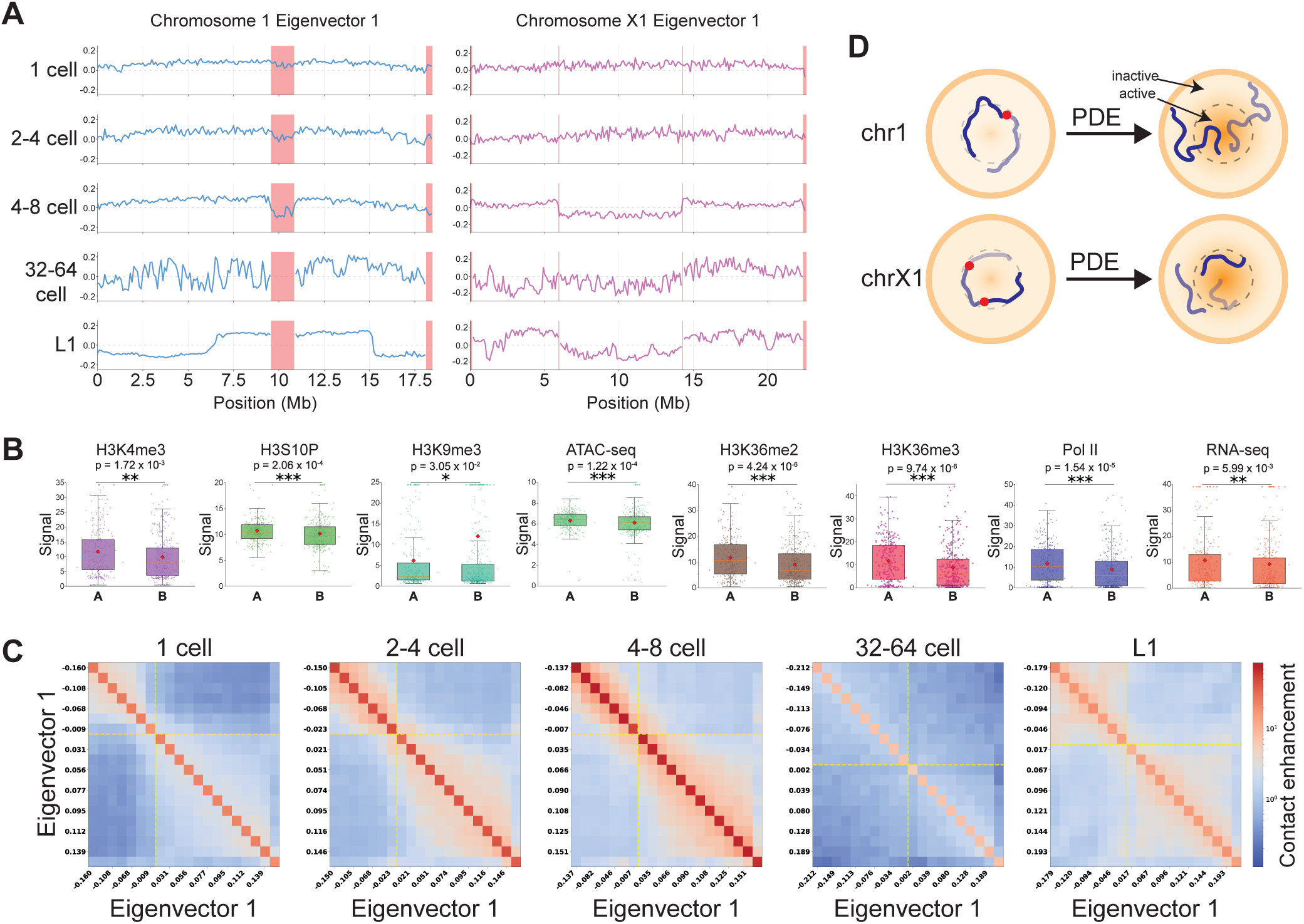
PDE splits chromosomes and causes 3D genome changes. **A.** Compartmentalization changes during *Ascaris* embryogenesis. Hi-C compartments were determined by using the eigenvector 1 values of chromosomes through the developmental stage. Eliminated regions are highlighted in red (see Figure S7 for compartments for all chromosomes). For chromosome 1, weak compartmentalization between its two future somatic derivatives is shown before PDE. Following PDE, the 3′ end of chromosome 1a interacts with the 5′ end of chromosome 1b within the active compartment, while the 5′ end of chromosome 1a and the 3′ end of chromosome 1b remain in the inactive compartment (see also Figure 5D). For chromosome X1, after PDE split it into three somatic chromosomes, the two terminal fragments occupy the same compartment while the central fragment is segregated away (see also Figure 5A). **B.** Multiomics analysis confirms the defined compartments. Active chromatin marks are enriched in A compartments, while a heterochromatin mark is enriched in B compartments. Each point represents a 100 kb genomic region. A two-sided Mann-Whitney U test was performed to calculate the statistical significance for each comparison (*** p<0.001, ** p<0.01, and * p<0.05). **C.** Dynamics of compartment interactions through development. A/B compartment interactions are shown using saddle plots. Genomic bins are sorted by PC1 value and binned into 20 quantiles (both axes). Contact enrichment within each bin pair is displayed (red = high; blue = low). Dashed yellow lines mark the A/B boundary (PC1 = 0). **D**. Model of chromosomal rearrangements following PDE. The germline chromosomes are not strongly compartmentalized in the early embryos. After PDE, the resulting somatic chromosomes reorganize their compartments. The two chromosomes were modeled based on data from Figure 5A.

### PDE-induced karyotype changes lead to compartmental change and somatic 3D genome reorganization

*Ascaris* PDE leads to dramatic karyotype changes and the splitting of 11 of the 24 germline chromosomes into 23 of the 36 somatic chromosomes [21] (the other 13 germline chromosomes do not have breaks in the middle of the chromosomes). To investigate whether and how large-scale chromosomal contact domains change after PDE, we first determined the A/B compartmentalization pattern using the Hi-C data (Figure 5A and Figure S7). This analysis designates each genomic region as either transcriptionally active (A compartment) or inactive (B compartment). Compartmentalization is weak during early development, up to the embryonic stages of PDE (4-8 cells), after which compartments are strengthened (Figure 5A and Figure S8A). This strengthening of compartments likely reflects both large-scale changes in karyotype, due to the increased number of chromosomes following PDE, and smaller-scale chromatin reorganization typical of early embryogenesis in nematodes [45, 46]. Overall, our analyses defined *Ascaris* A/B compartments and their changes during early embryogenesis.

We used a multi-omics approach to validate the compartments inferred from Hi-C (Figure 5B). Data from the 32-64-cell embryos (5 days) were used since they exhibit strong compartmentalization (Figure 5A) and comprehensive multi-omics data are available. Analysis of strongly compartmentalized regions (defined as EV1 > 0.15 or < -0.15) revealed that active chromatin marks, including H3K4me3, H3S10P, H3K36me2, and H3K36me3, are significantly enriched in A compartments. Conversely, H3K9me3, a marker of repressive chromatin, is significantly enriched in B compartments (Figure 5B). RNA-seq and ATAC-seq analyses also show significantly higher signal in A compartments, corroborating the compartment data. Furthermore, A compartments are enriched for active RNA Polymerase II (Figure 5B). Together, these analyses confirm that the defined compartments are consistent with transcriptional activities.

We observed notable compartmentalization changes surrounding the CBRs before and after PDE. Prior to PDE, the eliminated regions shifted into opposite compartments relative to retained regions. After PDE, chromosomes adopted distinct patterns of A/B compartmentalization as exemplified in the autosome chromosome 1 and sex chromosome X1 (Figure 5A and 5D). Interestingly, genome-wide analysis revealed that in the 32-64-cell embryos, a stage immediately following PDE, the Hi-C compartments exhibit the most variable structure (Figure 5A). This coincides with a significant drop in saddle strength that measures the degree of separation between active and inactive compartments (Figure 5C and Figure S8B). This suggests that the 32-64-cell stage is likely a transition phase for compartmentalization, where future somatic compartments are not fully established. This transition is further demonstrated by the compartment identity switching (A-to-B and B-to-A): it is stable before PDE (1-cell; Figure S8C) but shows a significant increase in A-to-B switching at 32-64-cell embryos and more B-to-A switching between the 32-64-cell and L1 stages. Notably, compartmentalization changes for the somatic chromosomes are stabilized at the L1 stage (Figure 5A and 5D), where the somatic program is firmly established. Overall, our findings indicate that PDE-induced karyotype changes contribute to the reorganization of the 3D genome in somatic cells.

### CBRs in *Parascaris* form transient and pairwise interactions during PDE

The related horse parasite *Parascaris* also undergoes PDE during 2-32 cell embryonic stages [7, 19]. *Parascaris* has a single germline chromosome with the majority of the DNA (>85%) in the two heterochromatic arms that will be eliminated. Seventy-two CBRs are interspersed in the middle of the euchromatic chromatin regions [19]. To determine if the CBRs in *Parascaris* also form 3D genome interactions, we performed Hi-C in embryos before (1 cell; 10 hr), during (2-4 cells; 17 hr), and after PDE (32-50 cells; 36hr). In 1-cell embryos before PDE, the whole chromosome is largely unstructured at the 3D genome level (Figure 6A and Figure S9). However, in the 2-4-cell stage, where PDE begins, we observed a striking pattern of CBR interactions: neighboring pairs of CBRs that will form the ends of future somatic chromosomes interact with each other (Figure 6A-B). When the Hi-C contact frequencies were normalized by the expected pattern of distance decay, 36 sites of higher-than-expected interactions became apparent that match the ends of the 36 future somatic chromosomes (Figure 6B). Like *Ascaris*, these CBR interactions are largely formed in the retained regions (Figure 6C). Quantification of these pairwise CBR interactions shows peak interactions at the 2-4 cell stage whereas these interactions are significantly reduced in 4-8 cell embryos after PDE (Figure 6D-E). Overall, these data show transient and specific 3D CBR interactions occur only during PDE, suggesting a potential role of the pairwise CBR interactions in *Parascaris* PDE (Figure 6F).

**Figure 6.**
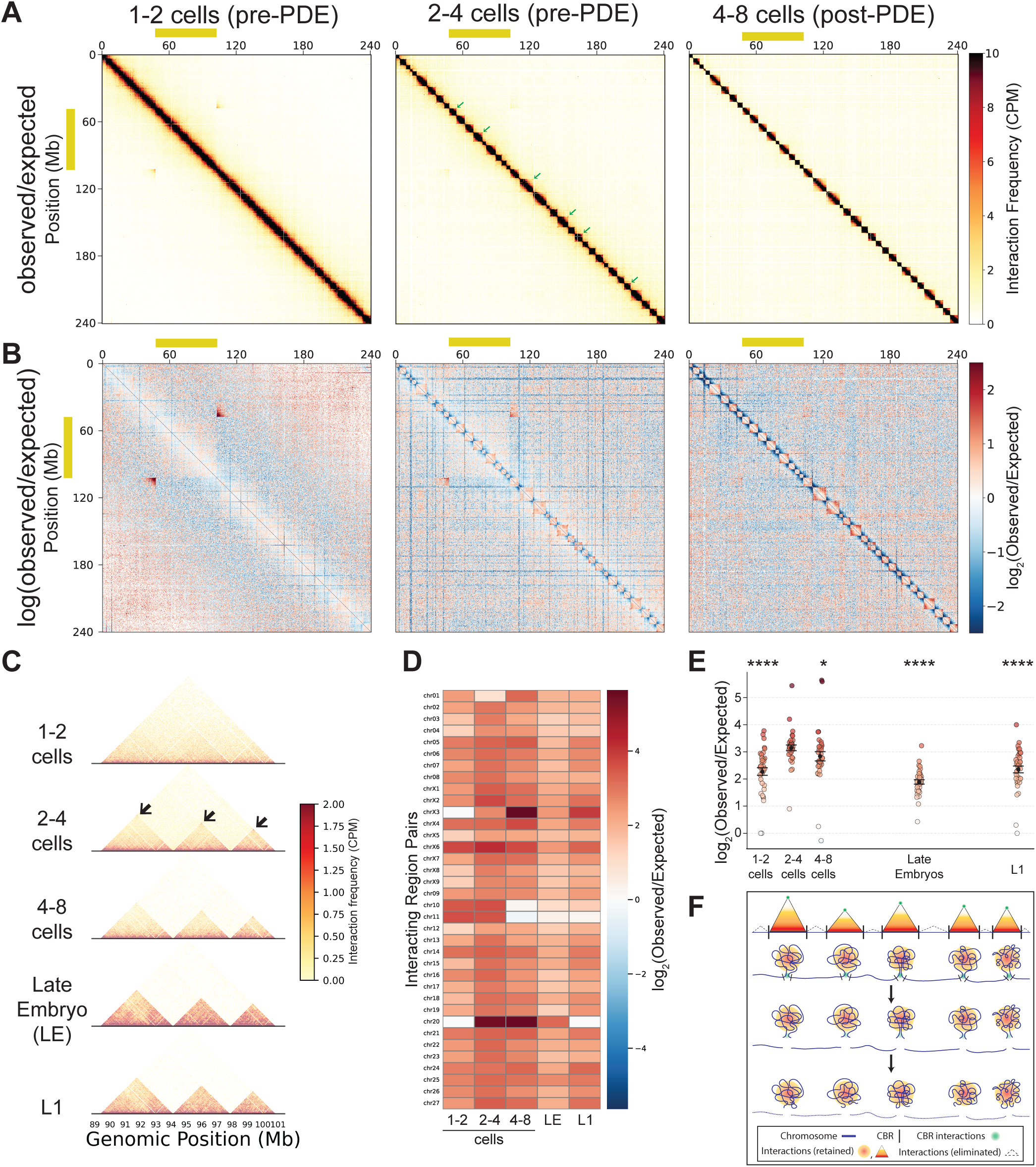
Transient end-to-end cis interactions between *Parascaris* CBRs during PDE. **A.** Hi-C contact map of the *Parascaris* embryos before and during PDE. The interaction between the boundaries of the sex chromosome region (yellow box) is due to the 25% of the Y chromosome in the population of embryos that lack the sex chromosome region (karyotype: female = XX and male = XY).[19] **B.** Pairwise CBR interactions during *Parascaris* PDE. Observed/expected contact ratio map for the same stages as in Figure 6A. Strong focal interactions are visible between the two CBRs that define the ends of each future somatic chromosome. **C.** Transient CBR interactions through *Parascaris* embryogenesis. Developmental Hi-C maps show that these end-to-end CBR interactions are not present in early embryos or prominent in late embryos. They peak at the 2-4-cell stage (black arrows), where PDE occurs. **D.** Heatmap showing enrichment of pairwise somatic chromosome ends in the germline genome during PDE. Each cell represents a single pair of coordinates based at somatic chromosome ends. Values are shown as log_2_(observed/expected). **E.** Plot of log2(observed/expected) values in D. Each dot represents a single region pair, colored by its log₂(O/E) value. Error bars are one standard error of the mean. A two-sided Mann-Whitney U test was performed to calculate the statistical significance for each comparison between the 2-4 cell sample and every other sample (**** p<0.0001 and * p<0.05). **F.** A model for CBR interactions during *Parascaris* PDE. Germline Hi-C interactions among future somatic chromosomes are shown with red-orange-yellow-while scale triangles, while interactions among eliminated regions are outlined with dashed lines (not to scale). Green spots represent interactions between CBR pairs near the ends of each future somatic chromosome. Note the resemblance of the pre-PDE chromosome structure to the “beads-on-a-string” model[69].

### Conserved compartment structure in *Parascaris* supports a role for PDE in somatic nuclear reorganization

Previous comparative analyses have shown that *Ascaris* and *Parascaris* have the same set of somatic chromosomes after PDE at sequence level [19]. To determine if similar programmed compartmentalization changes also occur in *Parascaris* at 3D genome level, we performed Hi-C on matching embryonic stages in *Parascaris*. Analysis of the compartments in *Parascaris* showed a strong correlation between the two species in post-PDE stages (Figure 7A). The majority of the matching somatic chromosome pairs (30/36; 83%) have a positive correlation coefficient of the eigenvector value ranging from 0.19 to 0.95 (median = 0.49) (Figure 7A). Importantly, for many matching pairs, such as chr01a/chr05, chr06b/chr07, chr02a/chr16, and chr14/chr21, the compartment patterns seen in *Ascaris* somatic chromosomes after PDE are mirrored in the somatic chromosomes of *Parascaris* (Figure 7A). Considering the divergence of the two species (over 10 million years [20]) and the sampling variations, the similarity of the Hi-C compartments is highly significant. Overall, our result suggests that PDE not only generates the same somatic chromosomes in these two parasites, but the resulting chromosomes undergo the same programmed compartmental changes after PDE, indicating a conserved somatic 3D genome after PDE.

**Figure 7.**
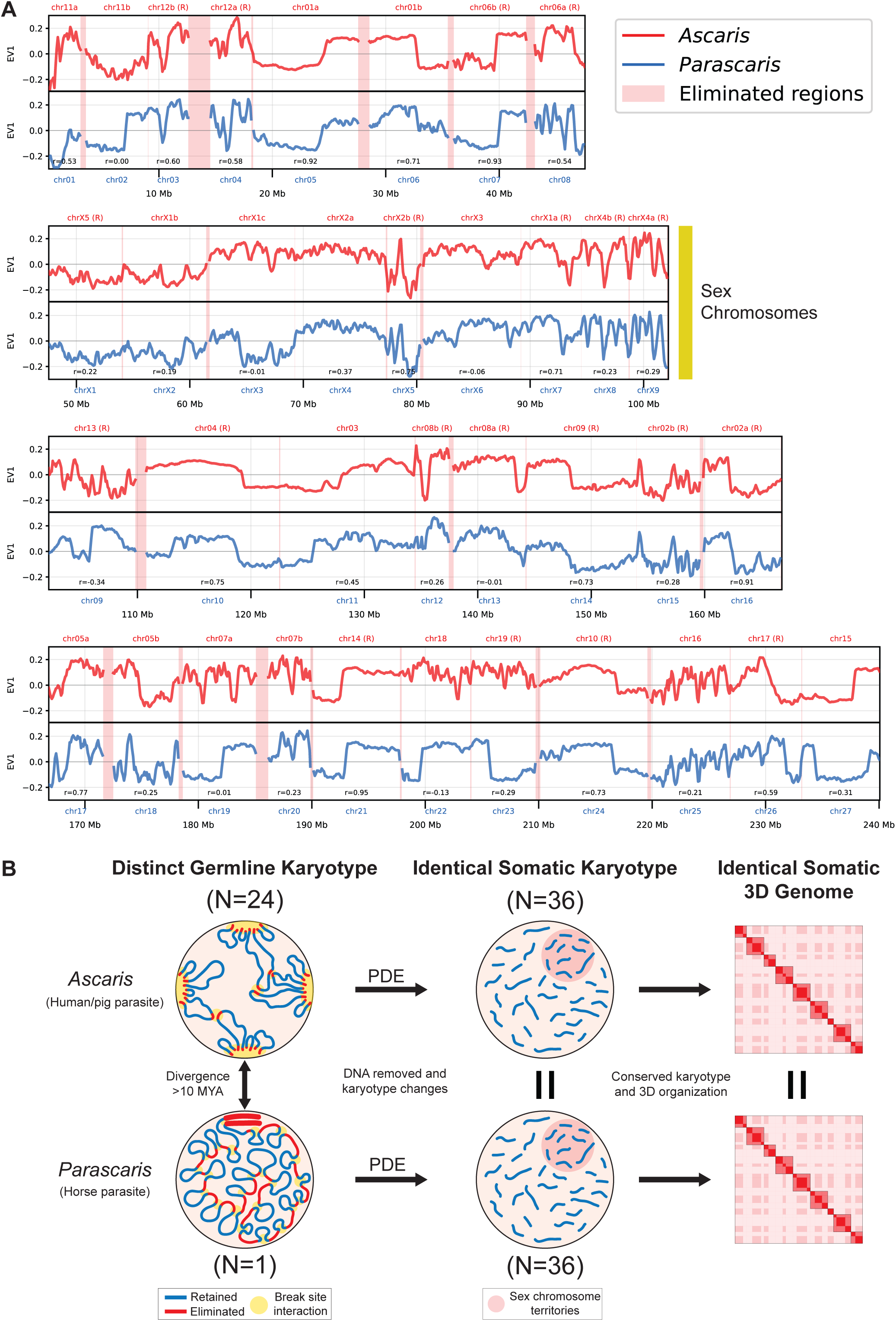
Chromosomal compartments are conserved between *Ascaris* and *Parascaris*. **A.** Eigenvector 1 values show conserved compartmentalization between *Ascaris* (red) and *Parascaris* (blue) in L1 larvae stages (after PDE). Positive values represent A (active) compartments and negative values represent B (inactive) compartments. Eliminated regions are shown with red highlights. The chromosomes are ordered based on their positions along the length of the single germline chromosome of *Parascaris*, with the coordinates shown at the bottom in Mb. Matching somatic chromosomes in *Ascaris* that are flipped on the x-axis (i.e., genome coordinates) are labeled with (R). **B.** PDE converts distinct germline karyotypes in *Ascaris* and *Parascaris* into identical somatic karyotypes with identical somatic 3D genome organization. See text in the Discussion section for a description of this summary figure.

## Discussion

In the parasitic nematode ascarids, DNA double-strand breaks (DSBs) occur within the chromosomal breakage regions (CBRs) of the germline genome, initiating programmed DNA elimination (PDE) (Figure 1A). This process results in a drastically reduced somatic genome. However, the mechanistic understanding of this process remains elusive. Here, using discrete staged embryos, we carried out Hi-C to examine the 3D genome organization before, during, and after PDE in both the *Ascaris* and *Parascaris* parasites. Two major findings emerged from our Hi-C analyses and comparison (summarized in Figure 7B). First, the 3D genome interactions among the CBRs are associated with PDE, indicating 3D genome may facilitate the break site recognition, DSB generation or DNA repair. Second, PDE lead to substantial and conserved compartmentalization changes in the somatic chromosomes of both nematodes, suggesting that by removing the 3D genome constraints caused by the to-be-eliminated DNA in the germline genome, PDE could play a critical role in reorganizing the somatic 3D genomes.

### Recognition of CBRs through 3D genome organization

*Ascaris* CBRs interact with each other prior to and during PDE (Figure 1). These interactions extend beyond the sites of CBRs, often covering the neighboring ∼200 kb of retained genome regions (Figure 3). Notably, many CBRs are located at the boundary of the spatially organized chromosome ends (Figure 3A), and significant insulation is observed in eliminated regions prior to elimination (Figure 3C). These findings suggest that CBRs are unique within *Ascaris*’s 3D genome organization and spatially demarcate the retained and eliminated sides of the DNA. Previous molecular data suggest CBR recognition is likely not dependent on specific sequences [21, 22]. Our Hi-C results lead us to propose a model for a sequence-independent mechanism of CBR recognition during *Ascaris* PDE (Figure 3E and Figure 7B). We hypothesize that CBRs interact with each other at foci where the machinery for DNA breaks and telomere healing is concentrated (PDE focus; Figure 3E and Figure 7B). The model does not require all CBRs to be at the same PDE focus, nor do these interactions need to occur simultaneously. The population-level Hi-C interactions observed among CBRs during PDE, which are lower than constant interactions but much higher than background-level interactions, support the model (Figure 1). The model may also provide a mechanism that doesn’t require nucleotide precision to generate DSBs [38], although it is plausible that other attributes of the CBRs and their localization could enhance the formation and function of PDE foci.

Chromosome end interactions that form the telomere bouquet are a common form of 3D chromosome organization during meiosis [47–50]. However, the interactions we observed at chromosome ends are unlikely caused by the telomere interactions. This is because the terminal CBRs have significantly higher trans chromosomal interactions at the retained sides of breaks, which are farther away from telomeres compared to the eliminated sides (Figure 3). Furthermore, we noticed that internal CBRs, which are far away from telomeres, also exhibit similar types of interactions (Fig. 1B-C). Consequently, we believe that the interactions observed at chromosome ends are likely unique to CBRs but not linked to telomeres. However, due to their locations near the ends of the chromosomes, we currently cannot determine whether the dynamic changes of interactions at the ends are caused by CBRs, telomeres, or both. Notably, we achieved a very high resolution of our Hi-C data by using multiple restriction digestions and high sequencing depth (for instance, 5 kb for Figure 3). Using these high-resolution Hi-C maps, we did not identify any other significant long-range or interchromosomal pairwise interactions (Figures 1, 2, 6). Thus, the CBR interactions stand out as a unique feature of the 3D genome in these nematodes, setting it apart from any other nematode species examined so far [33, 51–54]. The CBRs are the sites where DNA breaks occur, and evolutionarily, they are functionally conserved in both *Ascaris* and *Parascaris*. This highlights the significance of these interactions in PDE species.

A plausible mechanism for organizing the PDE foci is through phase-separated condensates [40, 55–58]. In other systems, such as olfactory receptor clustering in mammalian cells, phase-separated condensates can reinforce strong interchromosomal interactions similar to the ones observed between CBRs [59]. We suggest proteins and RNAs involved in DNA breaks and telomere addition may concentrate in PDE foci during DNA elimination. For instance, during *Ascaris* PDE, DSB ends are resected to generate long single-strand DNA (ssDNA) overhangs for telomere addition [38]. The ssDNA likely binds to RPA for protection and telomere synthesis [60–62]. The RPA protein complex is known for its strong propensity to form dynamic condensates [63]. Exogenous DSBs would not be recruited to these condensates, ensuring they are not healed with telomeres [38]. In addition, existing germline telomeres and shelterin components could also contribute to the formation of liquid-like droplets [64, 65]. This may explain why CBRs at the boundaries of spatially organized 3D domains are mostly observed at the ends of germline chromosomes where telomeres are present (Figure 3 and Figure S4). Furthermore, there is precedence for co-localization of chromatin regions at sites enriched for processing enzymes such as RNA splicing and nuclear speckles [66]. Additional factors, such as sequence- or nucleosome-dependent elements, may further contribute to the positioning of the CBRs for foci formation and DSB generation. Overall, our data suggest that the CBRs represent a unique genomic feature in nematode PDE at the boundary between the retained and eliminated DNA, and the 3D genome organization of the CBRs and chromosome ends may contribute to their recognition, foci formation, and the initiation and/or execution of PDE.

The CBR and chromosome end interactions precede PDE in the germline and early embryos (Figure 2). It remains to be determined if these interactions persist through fertilization and pronuclear fusion or if they are erased and re-established in the zygote. Even though these interactions exist in the germline, the factors associated with PDE foci formation and function may not be present, thereby preventing PDE from occurring. We speculate that these interactions in germline stages may serve a separate germline function, likely involved in chromosome organization during meiosis. In *C. elegans*, pairing and synapsis are achieved through the recognition of the pairing centers at one end of each chromosome by four related zinc finger proteins [67, 68]. Interestingly, these zinc finger protein genes and many other genes involved in *C. elegans* pairing are absent in *Ascaris* and *Parascaris* [19], suggesting that nematodes with PDE may use different mechanisms for pairing. Our data provide support that indeed, these germline chromosomes can interact with each other using their chromosome ends (Figure 2). These interactions may be an intrinsic feature associated with the sequences at the chromosome ends. During PDE, these sequences are removed, and the interactions among retained regions near the CBRs are abrogated. Therefore, PDE may offer a means to remove chromosome ends that function in the germline but are no longer required or may even be harmful in somatic cells. Interestingly, some of these CBRs show interactions along the entire length of multiple other chromosomes, leading to the formation of long stripe patterns in the Hi-C map (Figures 1 and 2). These long stripes could potentially signify an active searching, scanning, or looping mechanism. Future studies using single-cell and live imaging techniques are needed to unravel the dynamics and biological significance of these interactions.

The CBRs from *Parascaris* also interact during PDE. However, unlike *Ascaris* CBRs that form extensive interactions among the two CBR groups (Figure 1B), *Parascaris* CBRs engage in pairwise interactions mostly for CBR pairs from the future ends of each somatic chromosome (Figure 6). The *Parascaris* pairwise CBR interactions are weaker than *Ascaris* CBR interactions and occur transitionally only during PDE stages. They are reminiscent of the “beads-on-a-string” model proposed based on an early cytology study [69], where the “beads” represent retained and condensed future somatic chromosomes, whereas the “string” represents internally eliminated DNA associated with more loose chromatin during PDE (Figure 6F). Thus, our Hi-C data corroborate the cytologic observation that the CBRs in *Parascaris* are at the boundaries of spatially organized chromosomes that could separate the retained and eliminated DNA at the 3D level (Figure 6). The differences in CBR interactions observed in these two nematodes suggest possible variations in the PDE mechanisms, some of which may be attributed to the unique organization of the single homologous pair of the 2.5-Gb *Parascaris* germline chromosomes, where none of the CBRs are at the ends of the chromosomes, and even the most terminal CBRs are ∼1 Gb away from both chromosome ends [19].

### PDE provides a means to reorganize the somatic chromosomes

PDE resolves fused germline chromosomes (*Ascaris* = 24 and *Parascaris* = 1) into the same set of 36 somatic chromosomes [19] (Figure 7B). We previously suggested that the restoration of the somatic karyotype may confer biological functions [19]. Here, we demonstrate that the somatic 3D genome organization, as exemplified by the compartmentalization changes following PDE, is conserved between the two species (Figure 7A). While the impact of these compartmentalization changes on individual genes may be limited or negligible, the aggregate effect on numerous genes influenced by these changes could be substantial. Although the evolutionary trajectories leading to the current germline chromosomes are different [19], partitioning of the germline chromosomes leads to somatic chromosomes that retain the “memory” of the ancestral 3D genome organization. This underscores the important role of PDE in splitting the fused germline chromosomes.

Chromosome reorganization is a common response to DSBs [70–73]. Unlike most DSBs that are repaired using non-homologous end joining or homologous recombination, DSBs induced by PDE in nematodes are healed through telomere addition, accompanied by DNA loss [38, 74]. This telomere-mediated repair process may also contribute to chromosome reorganization by forming condensates surrounding the break sites. Interestingly, analysis of the Hi-C data across all 36 somatic chromosomes revealed consistent temporal changes in the 3D genome structure, showing the decreased insulation at and around the DSBs during PDE (Figure 4A). This change may be attributed to 1) the liberation of CBRs from their germline chromosome locations, 2) the DSB and telomere healing mechanisms, and 3) the formation of new heterochromatic regions at the somatic chromosome ends.

In *C. elegans* and other rhabditid nematodes, Hi-C compartments are also observed [33, 51–54]. These compartments exhibit a general profile for autosomes, where the active (A) compartments are in the middle of the chromosomes and inactive (B) compartments are in the arms of the chromosomes, forming a linear B-A-B profile along its arm-center-arm [75]. This B-A-B profile is consistent with expression data that more conserved and active genes are located in the middle of the chromosomes [76]. Comparative analysis indicates that this compartmental organization may be associated with the ancestral linkage groups in nematodes (known as Nigon elements [77, 78]), as fused chromosomes retain the centrality of the active compartment [79, 80]. Interestingly, this compartment profile is not consistently observed in *Ascaris* and *Parascaris* somatic chromosomes, despite that Nigon elements are largely maintained in their genomes [19]. Some of these differences could be attributed to the more discrete early embryo stages used in our study, although Hi-C data from a strictly somatic tissue (such as the intestine) also did not exhibit a consistent B-A-B compartment profile as in *C. elegans* [19]. Since *Ascaris* and *Parascaris* belong to the distant clade III of nematodes, it is plausible that their compartment profiles may have diverged from those of rhabditid nematodes (clade V), where Nigon elements were originally defined. Although many of the observed compartmental changes are specifically linked to the break sites associated with PDE (Figures 5 and 7A), we recognize that PDE occurs alongside other dynamic developmental process during embryogenesis, thus it is also plausible that some of these compartmental changes may be part of the developmental program.

In the ciliate *Tetrahymena thermophila*, distinct higher-order chromatin organization were observed in the germline micronucleus and somatic macronucleus [81], suggesting that the 3D genome may play important roles during the development of the macronucleus chromosomes during PDE. Similarly, a recent study in the ciliate *Oxytricha trifallax* also suggested widespread 3D genome reorganization before the onset of PDE [82]. Another recent genomic study in the ciliate *Paramecium aurelia* found that the ends of the germline chromosomes contain helitron repeats, forming a distinct genomic compartment that is eliminated during early somatic development [83]. These findings in ciliates are consistent with our observation that CBRs spatially demarcate retained and eliminated DNA in nematodes (Figure 3).

## Conclusions

PDE is an exception to the genome constancy rule. The mechanisms of PDE, particularly in multicellular organisms, remain largely unknown. The association of DNA break sites with 3D genome organization provides new insights into a possible mechanism. PDE creates drastic karyotype changes, often leading to the splitting of chromosomes. These changes in chromosomes are an integral part of development, and they offer a novel experimental system to investigate the impact of DSBs and karyotype changes on the reorganization of the 3D genome. As the number of known PDE species increases, including many amenable to genetic studies, further research is promising to unravel the functions of the 3D genome in the mechanisms and consequences of PDE.

## Acknowledgments

We thank Martin Neilsen for the *Parascaris* material and Bruce Bamber, Jeff Myers, and Routh Packing Co. for their support and hospitality in collecting *Ascaris* material. We thank the Wang lab members for helpful discussions. We also thank Brandon Estrem, Mariano Labrador, and Dick Davis for comments, edits, and critical reading of the manuscript. This work was supported by NIH grants to J. W. (R01AI155588 and R01GM151551) and R.P.M. (R35GM133557) and the University of Tennessee Knoxville Startup Funds to J.W.

## Author contributions

J.R.S. and J.W. designed the project; J.R.S. made Hi-C libraries; J.R.S. improved the *Ascaris* genome assembly; J.R.S., T.X., and R.P.M. performed Hi-C data analysis; J.R.S. wrote the initial draft; J.W. wrote the manuscript; J.R.S., R.P.M., and J.W. reviewed and edited the manuscript; J.W. and R.P.M. provided supervision; and J.W. managed the project and provided funding.

## Materials and Methods

### Nematode Sample Collection and Staging

*Ascaris* ovaries and testes were dissected from adult worms. *Ascaris* 1-cell embryos were collected from fresh ovaries by dissolving the tissue with two treatments: 0.5 N NaOH for 60 min at room temperature, followed by washing in cold PBS buffer, pH 2.0. Embryonation of *Ascaris* zygotes was performed in PBS, pH 2.0, at 30°C while shaking for 10 hr (1 cell), 48 hr (2-4 cells), 60 hr (4-8 cells), 5 days (32-64 cells), and 10 days (L1 larvae). Ovary, testes, and staged embryos were frozen in liquid nitrogen and stored at -80°C until use.

Collection of *P. univalens* zygotes and embryonation were performed as previously described[19]. Dissected zygotes were stored at 4°C. Embryonation was performed in PBS, pH 2.0, at 37°C while shaking for 0 hr (1 cell), 17 hr (2-4 cells), 24 hr (4-8 cells), 48 hr (100-500 cells), 60 hr (300-800 cells), or 72 hr (L1 larvae).

Decoating of *Ascaris* and *Parascaris* embryos was performed by resuspending embryos in 0.4 N KOH/1.4% sodium hypochlorite and incubating for 90 min at 30°C, shaking at 100 RPM. Embryos were washed in cold PBS, pH=7.0, five times and pelleted at 1,250 x g for 5 min to estimate the volume of decoated embryos. Staging and decoating of *Ascaris* and *Parascaris* embryos were confirmed by visual inspection under a compound light microscope at 40X magnification.

### Hi-C Library Preparation

Fixation was performed using the Arima-HiC 2.0 kit standard input protocol for *Ascaris* ovaries and testis. For both *Ascaris* and *Parascaris* embryos, the low input protocol was used, with the following modification: cell membranes were disrupted by forcing cells through a small metal dounce in 5 mL of PBS + TC buffer (22% formaldehyde, 50 mM HEPES pH 8.0, 0.5 mM EGTA, 1 mM EDTA, 100 mM NaCl) kept on ice. Hi-C was performed using the Arima-HiC 2.0 kit according to the manufacturer’s protocol (Arima, Cat#A410110). After cross-linking and digestion, samples were fragmented using the Covaris M220 ultrasonicator (Covaris, Cat#500295) with the following settings: temperature = 7°C, peak incident power = 75 W, duty factor = 5%, cycles per burst = 200, time = 70 s. This generated fragments of around 500 bp. Size selection was performed using AMPure XP magnetic beads (Beckman Coulter, A63881).

Illumina libraries were constructed using the Swift Biosciences Accel-NGS 2S Plus DNA Library kit (Swift Cat# 21024) and the NEBNext Ultra II DNA Library Prep Kit (NEB). Libraries were amplified using the KAPA Library Amplification kit (Roche Cat# KK2620) according to the manufacturer’s protocols. Sample concentrations were measured with a DNA Qubit (Invitrogen, Cat# Q33238), and library fragment sizes were confirmed using a TapeStation 4200 (Agilent). Illumina libraries at 2 x 150 bp were sequenced using a NovaSeq 6000 at the University of Colorado Anschutz Medical Campus Genomics Core to a depth of 16-60X.

### Multi-omics Data Generation

The following datasets used for multi-omics analysis were previously published: ATAC-seq, RNA-seq, ChIP-seq for histone marks (H3K4me3, H3S10P, H3K36me2, H3K36me3, H3K9me3), and RNA Polymerase II[21, 22, 84, 85].

### Computational Analysis

#### *Ascaris* Genome Assembly and Improvement

The germline *Ascaris* genome assembly was systematically refined through iterative manual curation guided by Hi-C contact maps. Initial improvements included correcting obvious mis-assemblies, filtering unplaced scaffolds, and repositioning genomic segments based on artifactual Hi-C interaction patterns. Throughout this process, Hi-C contact heatmaps were generated at multiple resolutions to visually identify mis-assemblies. Corrections were implemented iteratively until contact maps displayed canonical chromatin interaction patterns with minimal obvious structural artifacts. Previously unplaced contigs were integrated into chromosomes where Hi-C linkage data supported their placement. The assembly was further improved by breaking the genome at scaffold junctions and re-scaffolding using Hi-C proximity ligation data. Chromosome termini were extended toward telomeres by mapping PacBio long reads to the corrected assembly, filtering previously incorporated reads, and using unmapped telomere-containing reads and contigs to extend chromosome ends. CBR coordinates from the previous *Ascaris* assembly were transferred to the updated genome using BLAST[86] to define both terminal and internal CBR sites in the updated assembly.

#### Hi-C Data Processing

Paired-end Hi-C reads were aligned to the *Ascaris* reference genome using Bowtie2[87], and contact matrices were generated and iteratively corrected with HiC-Pro[88, 89]. AllValidPairs files were converted to .hic format using hicpro2juicebox from HiC-Pro for downstream analysis. Standard HiC-Pro quality control metrics were computed for all samples, including mapping rates, valid interaction percentages, and trans/cis interaction ratios. Repetitive sequences were not masked in the primary analysis; a control analysis with repeat masking was performed separately to assess the impact of repeat filtering (Figure S2). For contact map visualization, allValidPairs files were converted to .hic[90] and .cool/.mcool formats[91]. Genomic heatmaps were generated at 250 kb resolution using raw contact counts normalized to counts per million (CPM): CPM = (raw counts / total valid pairs) × 10⁶. Downstream analyses used both .hic files and HiC-Pro-generated contact matrices at appropriate resolutions for each application.

#### Region-Based Contact Matrix Construction

Genomic bins were mapped to break regions annotated as terminal or internal. Region-level contact matrices were constructed by aggregating all bin-level interactions between region pairs. Contact frequencies were normalized to counts per million (CPM), and intra-chromosomal (cis) contacts were masked.

#### Principal Component Analysis of Interaction Profiles

Principal component analysis was performed on trans-chromosomal interaction profiles to assess global organizational differences between terminal and internal regions. Each region was represented by a feature vector containing its normalized contact intensities with all trans-chromosomal partners. 95% confidence ellipses around each region type cluster were calculated using a chi-square distribution.

#### Trans-Chromosomal Interaction Analysis at CBRs

Trans-chromosomal Hi-C interactions were quantified from allValidPairs files for pre- and post-elimination developmental stages. The genome was partitioned into 5 kb bins, and trans interaction counts (interactions between different chromosomes) were aggregated for each bin. Counts were normalized to counts per million (CPM) mapped reads. For each CBR, interaction profiles were computed by extracting bins within 500 kb windows upstream (into the retained region) to 100 kb downstream (into the eliminated region) around the CBR midpoint. Individual CBR profiles were aggregated across either all end CBRs or across all internal CBRs to generate meta-profiles for each developmental stage. To quantify which chromosomes contributed to the elevated trans interactions near the CBRs, partner chromosome contributions were represented proportionally based on normalized interaction frequencies.

#### Insulation Score Analysis

Chromatin insulation scores (where lower values indicate stronger insulation and higher values indicate weaker insulation) were calculated from Hi-C contact matrices at 20 kb resolution using a 100 kb insulation square size and 80 kb insulation delta span using FAN-C software[92]. To visualize insulation dynamics across programmed DNA elimination, insulation scores from pre-elimination samples were plotted directly on germline genome coordinates, while post-elimination samples were mapped back to pre-elimination coordinates using syntenic region mappings. Eliminated regions were highlighted but excluded from post-elimination plots to avoid artifacts. For genome-wide visualization, insulation scores were plotted for all chromosomes at each developmental stage (1 cell, 2-4 cells, 4-8 cells, 32-64 cells, L1 larvae). Smoothing was applied using a moving average filter (10 bin windows).

### Multi-Omics Analysis of New Somatic Chromosome Ends

To characterize the chromatin state at newly formed somatic chromosome ends following DNA elimination, we defined 100 kb terminal regions reaching from each CBR into the neighboring retained region. CBRs were separated for analysis by their presence near the end of chromosomes or internally/near the middle of a chromosome. For each developmental time point, multiple genomic and epigenomic datasets were analyzed as available: insulation scores (chromatin domain organization), RNA-seq (gene expression at 50 kb resolution), ATAC-seq (chromatin accessibility at 50 kb resolution), H3K9me3 ChIP-seq (heterochromatin mark at 50 kb resolution), and H3K4me3 ChIP-seq (active chromatin mark at 50 kb resolution).

For each 100 kb end or middle region, mean values were calculated by averaging all bins that overlapped the region. When multiple biological replicates were available (ATAC-seq, ChIP-seq), values were first averaged across replicates before regional quantification. Data were visualized across five developmental stages spanning pre-elimination (1 cell, 2-4 cells, 4-8 cells) through post-elimination (32-64 cells, L1 larvae) timepoints. Statistical comparisons between each time point were performed using two-sided Mann-Whitney U tests. Data were plotted as mean ± one standard error of the mean (SEM), with individual data points representing a single chromosome end. Sample sizes (n) represent the number of chromosomal regions analyzed at each time point.

#### A/B Compartment Analysis

To determine the compartments, we use Principal Component Analysis (PCA) on the normalized Hi-C contact matrix to identify the leading eigenvector. This primary PC is often reliable in identifying the A/B compartments[93]. However, for the *Parascaris* Hi-C analysis, the A and B compartments may need to be swapped in certain cases based on additional genomic data, such as histone markers, RNA-seq, and RNA pol II ChIP-seq (Figure S7). Additionally, in early embryos, the first PC may not accurately represent the A/B compartments due to limited distinct Hi-C interactions, among other factors. In such cases, we manually examine these instances and adjust the compartment call when necessary (Figure S7).

#### Validation of Compartment Identity with Multi-Omics Data

To validate the biological accuracy of A/B compartment assignments, we integrated genomic and epigenomic datasets at the 32-64-cell stage. This stage was chosen as it represented the most complete dataset and showed strong compartmentalization. Eigenvector values (100 kb resolution) were matched with multiple 10 kb resolution datasets, including RNA-seq, PRO-seq, ATAC-seq, and ChIP-seq for histone modifications (H3K4me3, H3K27me3, H3K9me3, H3K36me3, H3K36me2, H3K27me1, H4K20me1, H3S10p, and H2AK119ub), RNA Polymerase II, and centromeric proteins (CENP-A, CENP-C). For each 100 kb eigenvector bin, overlapping 10 kb omics bins were identified, and signal values were averaged to match resolutions. Bins were classified as A compartment (eigenvector > 0) or B compartment (eigenvector < 0). Statistical significance of signal differences between A and B compartments was assessed using two-sided Mann-Whitney U tests.

#### Compartment Strength Analysis

A/B compartment interactions were analyzed using saddle plots, which aggregate Hi-C contact frequencies as a function of compartment identity. For each developmental stage, 100 kb resolution contact matrices and eigenvector (PC1) values were used. Genomic bins were sorted by PC1 value and partitioned into 20 quantiles to ensure even distribution. Contact frequencies between all bin pairs were then aggregated according to their quantile assignments, creating a 20×20 saddle matrix. Saddle strength was quantified as the ratio of within-compartment to between-compartment interactions: (AA + BB) / (AB + BA), where AA represents mean contact enrichment between A compartment bins, BB between B compartment bins, and AB between A and B compartment bins. This measures compartment segregation, with higher values indicating stronger separation between euchromatic (A) and heterochromatic (B) compartments. Compartment strength was calculated as the mean absolute PC1 value. Compartment switching between consecutive developmental stages was assessed by comparing PC1 signs for each genomic bin. Bins were classified as stable (maintaining A or B identity), A-to-B switches, or B-to-A switches. Only bins with |PC1| > 0.005 were included in the switching analysis to exclude weakly defined compartments. Statistical significance of switching frequencies was evaluated using two-sided Mann-Whitney U tests.

#### Expected Contact Frequency Analysis

Expected Hi-C contact frequencies as a function of genomic distance were computed from .hic files using FAN-C[92]. For each sample, genome-wide expected contact matrices were generated at 100 kb resolution. Genome-wide expected contact profiles were combined across all samples by aligning contacts at equivalent genomic distances. Plots were normalized by scaling each sample’s profile such that the contact frequency at the first distance bin equaled the average across all samples at that distance, allowing direct comparison of decay rates independent of absolute interaction counts.

#### Chromosome Interaction Clustering Analysis

Whole-chromosome trans interaction patterns were analyzed to identify chromosomes with similar interaction profiles before and after programmed DNA elimination. Inter-chromosomal interactions were extracted from allValidPairs files for pre-elimination samples (24 intact chromosomes) and post-elimination samples (36 chromosomal segments following DNA elimination). Mitochondrial and unplaced scaffolds were excluded from analysis.

For each sample, raw interaction counts between chromosome pairs were compiled into symmetric interaction matrices by averaging reciprocal interactions (i.e., interactions from chromosome A to B and B to A). Self-interactions were excluded (set to NaN). Matrices were normalized using an expected-trans interaction model where expected interaction frequencies between chromosome pairs were calculated based on the total trans interaction counts for each chromosome. Normalized values were computed as observed/expected ratios. Interaction matrices were visualized as clustered heatmaps with hierarchical dendrograms displayed along both axes to illustrate chromosome relationships.

#### *Parascaris* Chromosome End Interaction Analysis

To quantify interactions between regions destined to become somatic chromosome ends, pairs of 100 kb windows centered on the terminal regions of each of the 36 somatic chromosomes were defined in germline genome. Interaction scores between each region pair were computed from ICE-normalized Hi-C contact matrices at 20 kb resolution. For each region pair, the mean contact frequency was calculated across all non-zero bin-pairs within the submatrix. Scores were normalized using an observed/expected approach: expected contact frequencies were computed as the mean contact frequency at each genomic distance, calculated within each chromosome to account for the single germline chromosome in pre-PDE samples and the 36 somatic chromosomes in post-PDE samples. Interaction scores were expressed as log₂(observed/expected) and visualized as heatmaps across developmental timepoints and as summary plots showing mean and median log₂(O/E) ± one standard deviation across all 36 region pairs.

#### Comparative Compartment Analysis Between Species

A/B compartment structure was determined for *Ascaris* and *Parascaris* from Hi-C data using principal component analysis (PCA) on normalized contact matrices at 100 kb resolution, implemented in FAN-C[92]. The first eigenvector (EV1) of the observed/expected contact matrix was computed genome-wide and oriented by correlation with GC content, where positive values indicate GC-rich, open A compartments and negative values indicate GC-poor, closed B compartments. Post-elimination *Parascaris* chromosomal segments were mapped back to pre-elimination germline coordinates to establish a common reference frame. *Ascaris* chromosomes were similarly mapped to *Parascaris* homologs using chromosome-level synteny[19], with coordinates scaled proportionally to account for length differences between species.

For each species, EV1 profiles were corrected for arbitrary sign by applying chromosome-specific polarity adjustments to ensure biological consistency across homologous regions. Profiles were smoothed using a rolling mean (window size = 3 bins) to reduce noise while preserving large-scale compartment structure. Eliminated regions were highlighted but excluded from quantitative analysis. The degree of compartment conservation between homologous chromosomes was quantified using Pearson correlation coefficients. For each chromosome pair, EV1 values from *Ascaris* and *Parascaris* were interpolated to 100 common genomic positions spanning the homologous region, and correlation was calculated after applying appropriate coordinate transformations (i.e., reversals for inverted syntenic blocks).

#### Computational Implementation and Artificial Intelligence Usage

Hi-C data processing and visualization were performed with custom Python and shell scripts. All analyses were implemented in Python 3.9 using NumPy v1.21.0 for linear algebra operations, pandas v1.3.0 for data manipulation, matplotlib v3.5.0 and seaborn v0.11.0 for visualization, and SciPy v1.9.0 for statistical tests. Script development and debugging were assisted by Claude Sonnet 4.1 and 4.5 and Opus 4.6 (Anthropic), with all outputs verified through manual inspection and comparison with published methods. Claude Sonnet 4.5 was used to refine technical descriptions for clarity and concision. All data and scripts are available in https://github.com/jsimmo45/Nematode-3D-Genome-PDE.

## Availability of data and materials

The datasets supporting the conclusions of this article are available in the NCBI GEO repository, GSE314626 and GSE315650, in https://www.ncbi.nlm.nih.gov/geo/query/acc.cgi?acc=GSE314626 and https://www.ncbi.nlm.nih.gov/geo/query/acc.cgi?acc=GSE315650. The datasets supporting the conclusions of this article are included within the article and its additional supplemental files.

