## Supplemental Figures for "Spatial genome organization in nematodes with programmed DNA elimination"

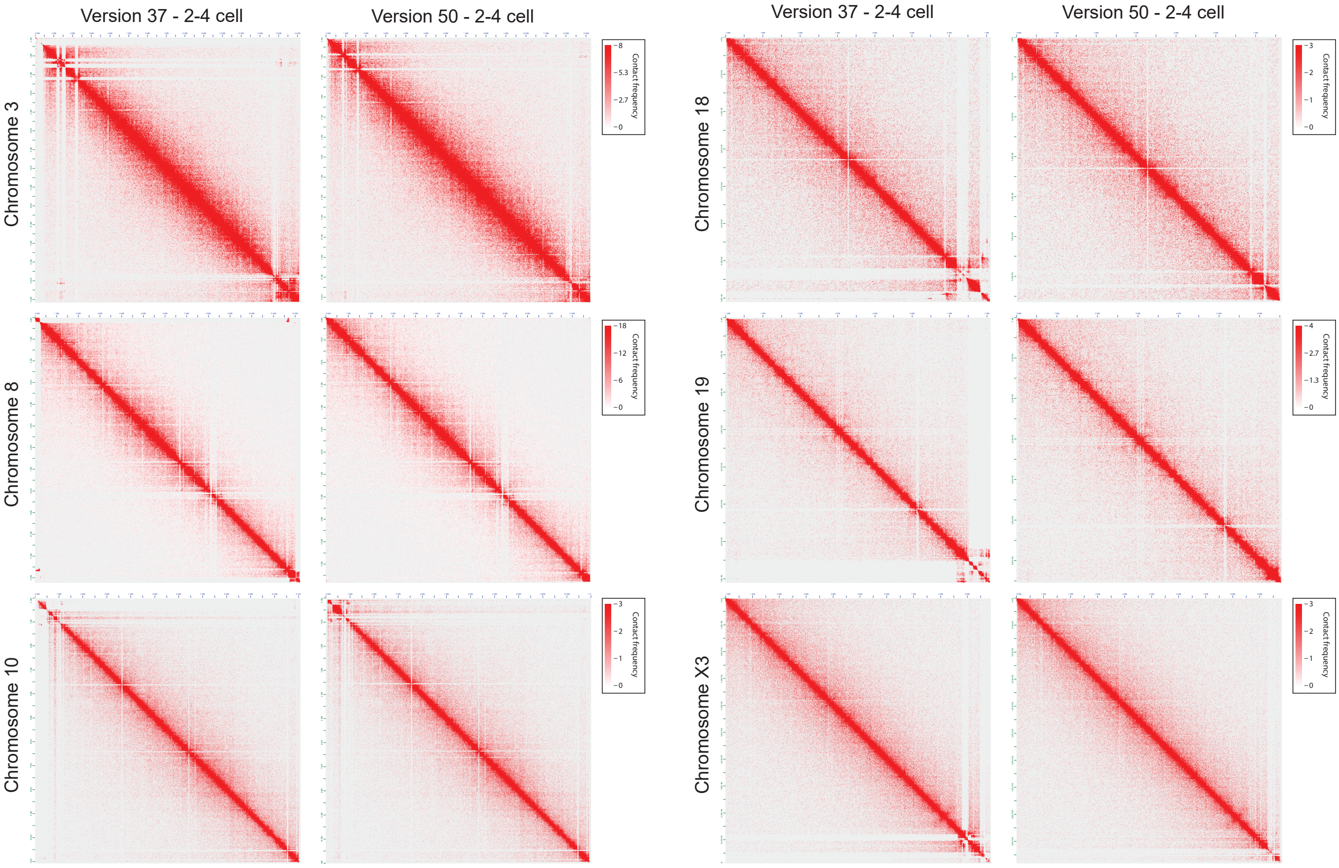

Version 37 - 2-4 cell
Version 50 - 2-4 cell
Version 37 - 2-4 cell
Version 50 - 2-4 cell
Chromosome 3
Chromosome 8
Chromosome 10
Chromosome 18
Chromosome 19
Chromosome X3
Contact frequency


### Figure S1

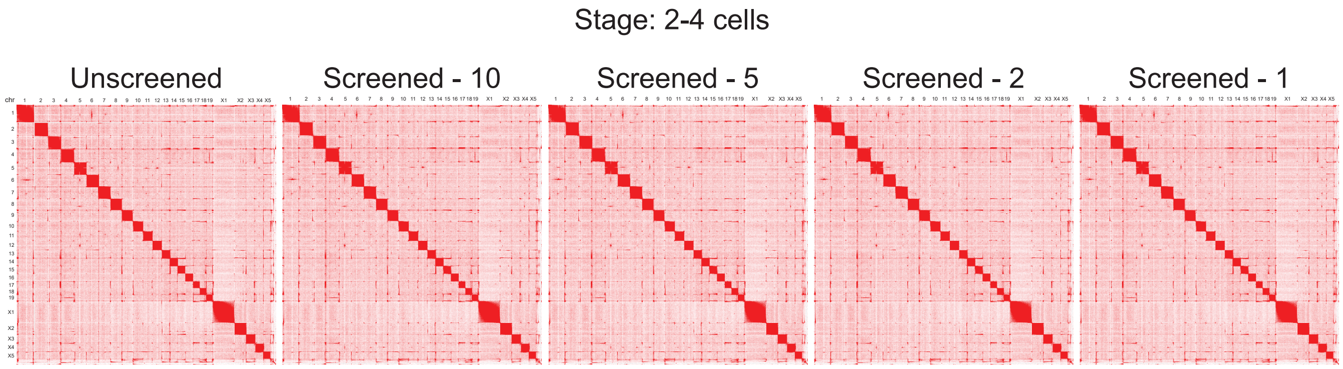

Stage: 2-4 cells
Unscreened
Screened - 10
Screened - 5
Screened - 2
Screened - 1


**Figure S2**

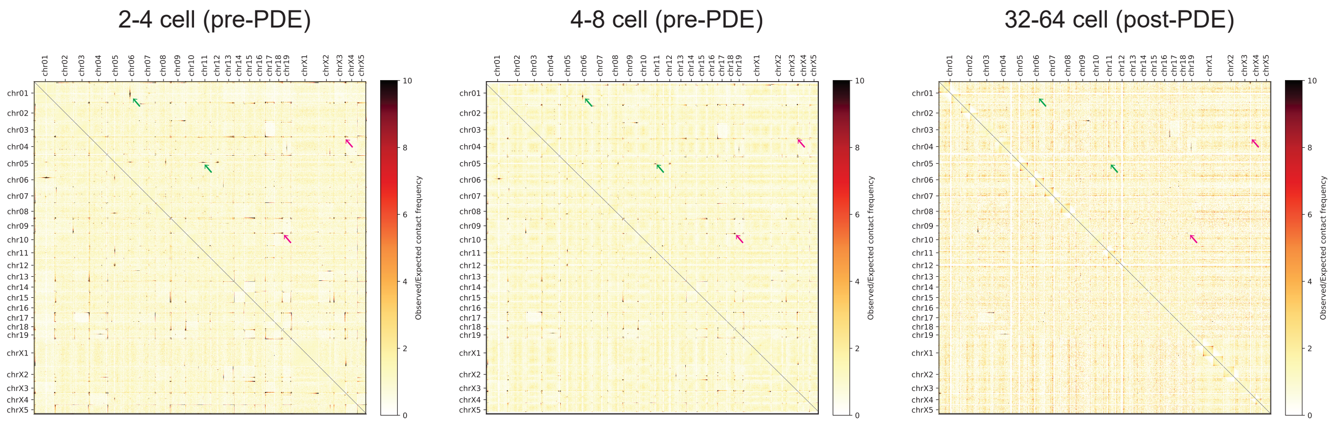
2-4 cell (pre-PDE)
4-8 cell (pre-PDE)
32-64 cell (post-PDE)
chr01
chr02
chr03
chr04
chr05
chr06
chr07
chr08
chr09
chr10
chr11
chr12
chr13
chr14
chr15
chr16
chr17
chr18
chr19
chrX1
chrX2
chrX3
chrX4
chrX5
Observed/Expected contact frequency
0
2
4
6
8
10

### Figure S3

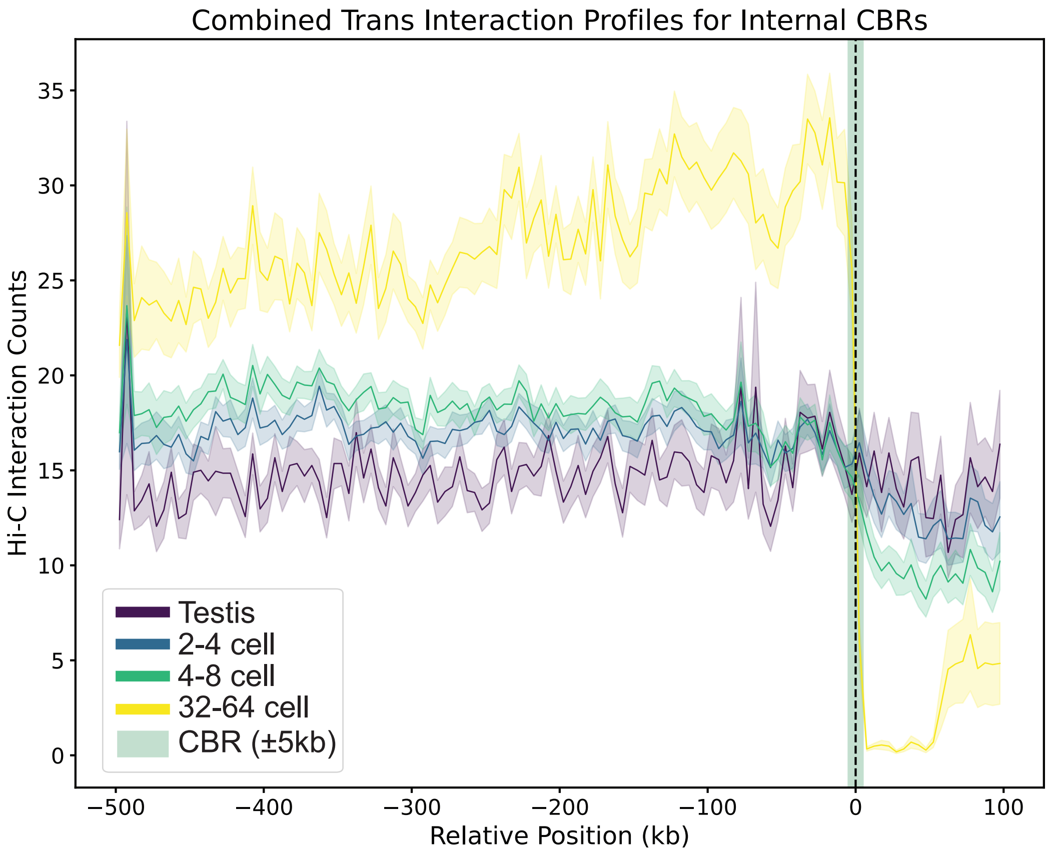
Combined Trans Interaction Profiles for Internal CBRs
Hi-C Interaction Counts
0
5
10
15
20
25
30
35
−500
−400
−300
−200
−100
0
100
Relative Position (kb)
Testis
2-4 cell
4-8 cell
32-64 cell
CBR (±5kb)

**Figure S4**

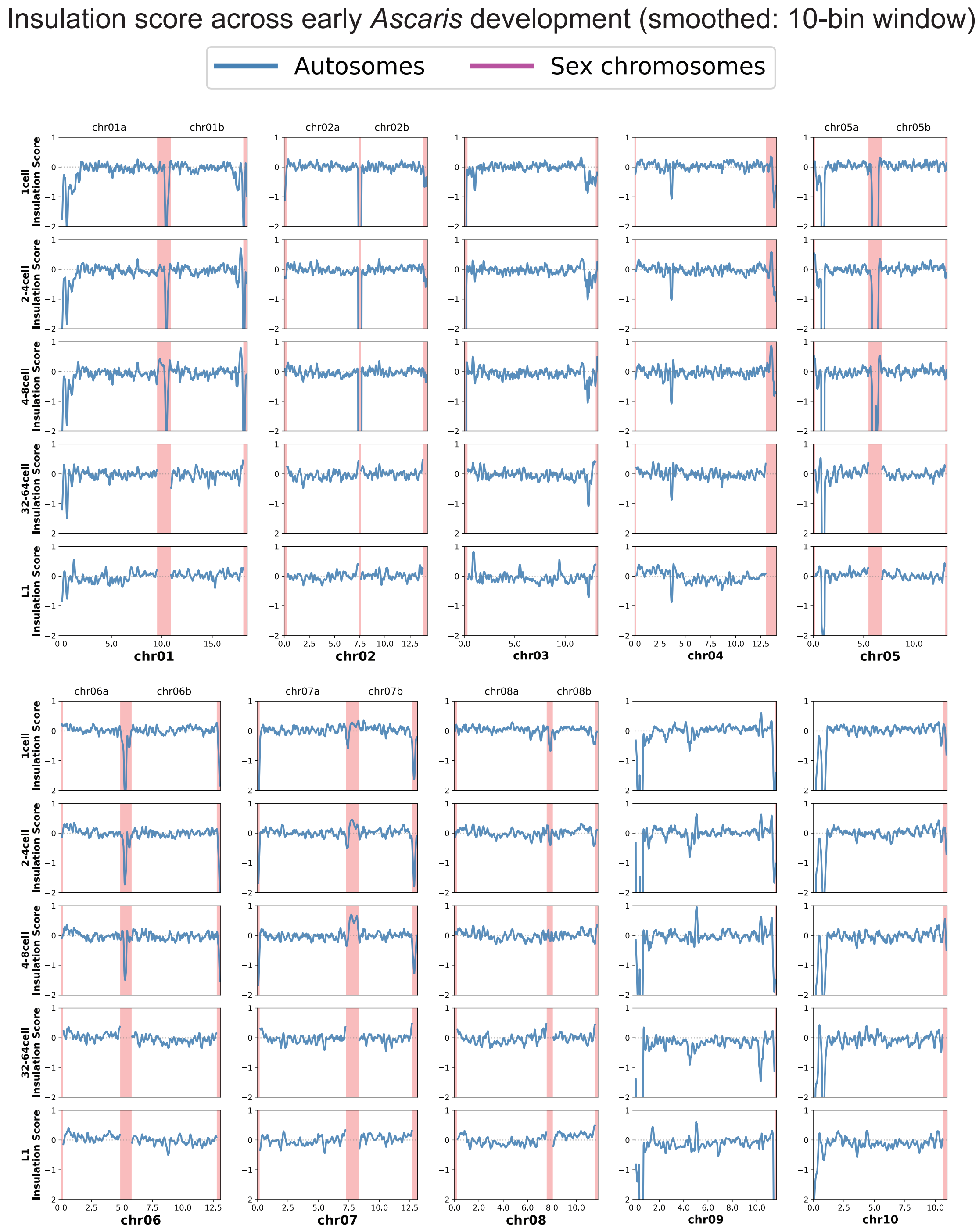
Insulation score across early Ascaris development (smoothed: 10-bin window)
Autosomes
Sex chromosomes
chr01a
chr01b
chr02a
chr02b
chr05a
chr05b
1cell Insulation Score
2-4cell Insulation Score
4-8cell Insulation Score
32-64cell Insulation Score
L1 Insulation Score
chr01
chr02
chr03
chr04
chr05
chr06a
chr06b
chr07a
chr07b
chr08a
chr08b
chr06
chr07
chr08
chr09
chr10

**Figure S5**

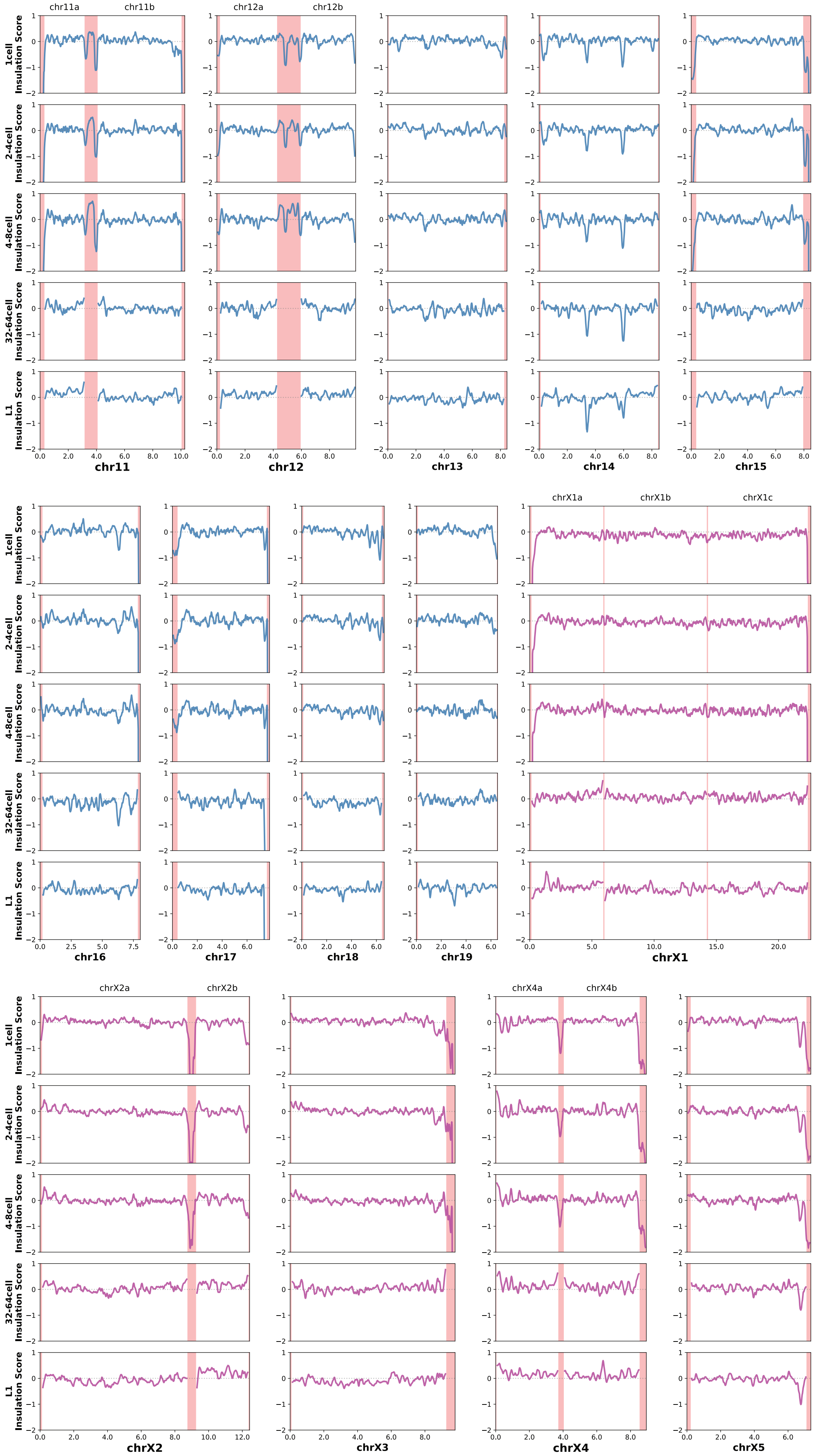


**Figure S5 cont.**

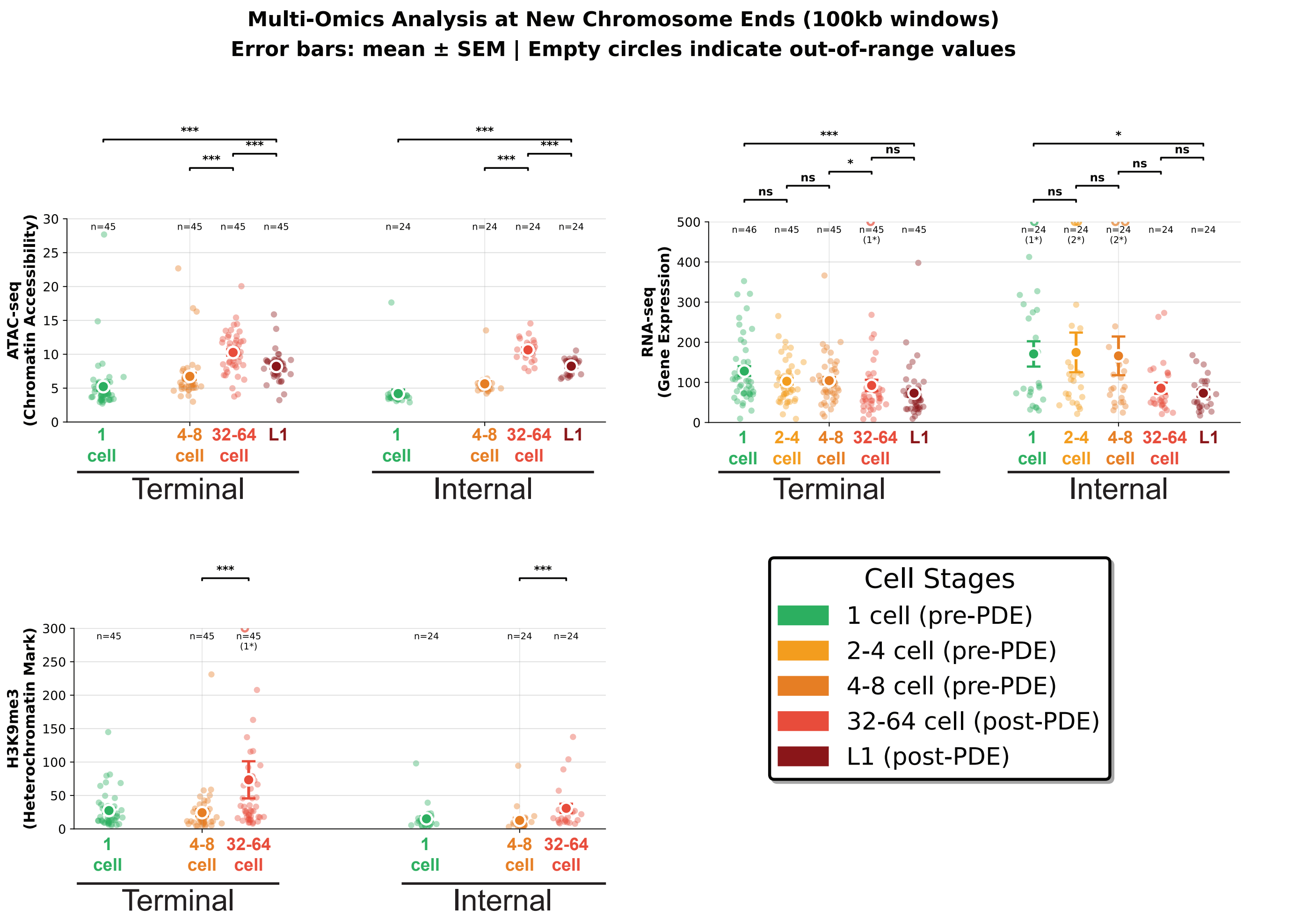
Multi-Omics Analysis at New Chromosome Ends (100kb windows)
Error bars: mean ± SEM | Empty circles indicate out-of-range values
ATAC-seq
(Chromatin Accessibility)
RNA-seq
(Gene Expression)
H3K9me3
(Heterochromatin Mark)
1 cell
2-4 cell
4-8 cell
32-64 cell
L1
Terminal
Internal
n=45
n=24
n=46
(1*)
(2*)
***
*
ns
Cell Stages
1 cell (pre-PDE)
2-4 cell (pre-PDE)
4-8 cell (pre-PDE)
32-64 cell (post-PDE)
L1 (post-PDE)

#### Figure S6

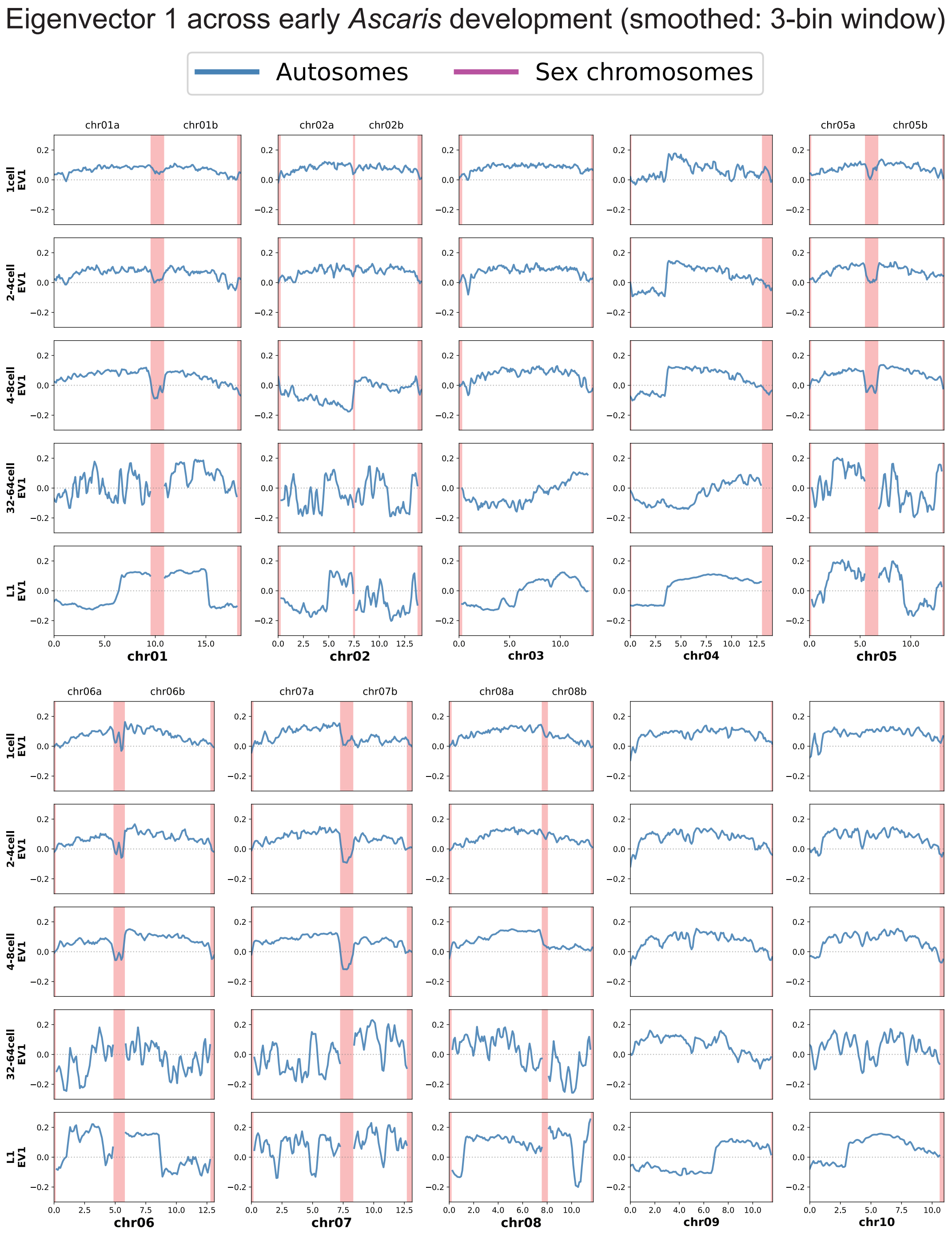
Eigenvector 1 across early Ascaris development (smoothed: 3-bin window)
Autosomes
Sex chromosomes
chr01a
chr01b
chr02a
chr02b
chr05a
chr05b
chr06a
chr06b
chr07a
chr07b
chr08a
chr08b
1cell EV1
2-4cell EV1
4-8cell EV1
32-64cell EV1
L1 EV1
chr01
chr02
chr03
chr04
chr05
chr06
chr07
chr08
chr09
chr10

**Figure S7**

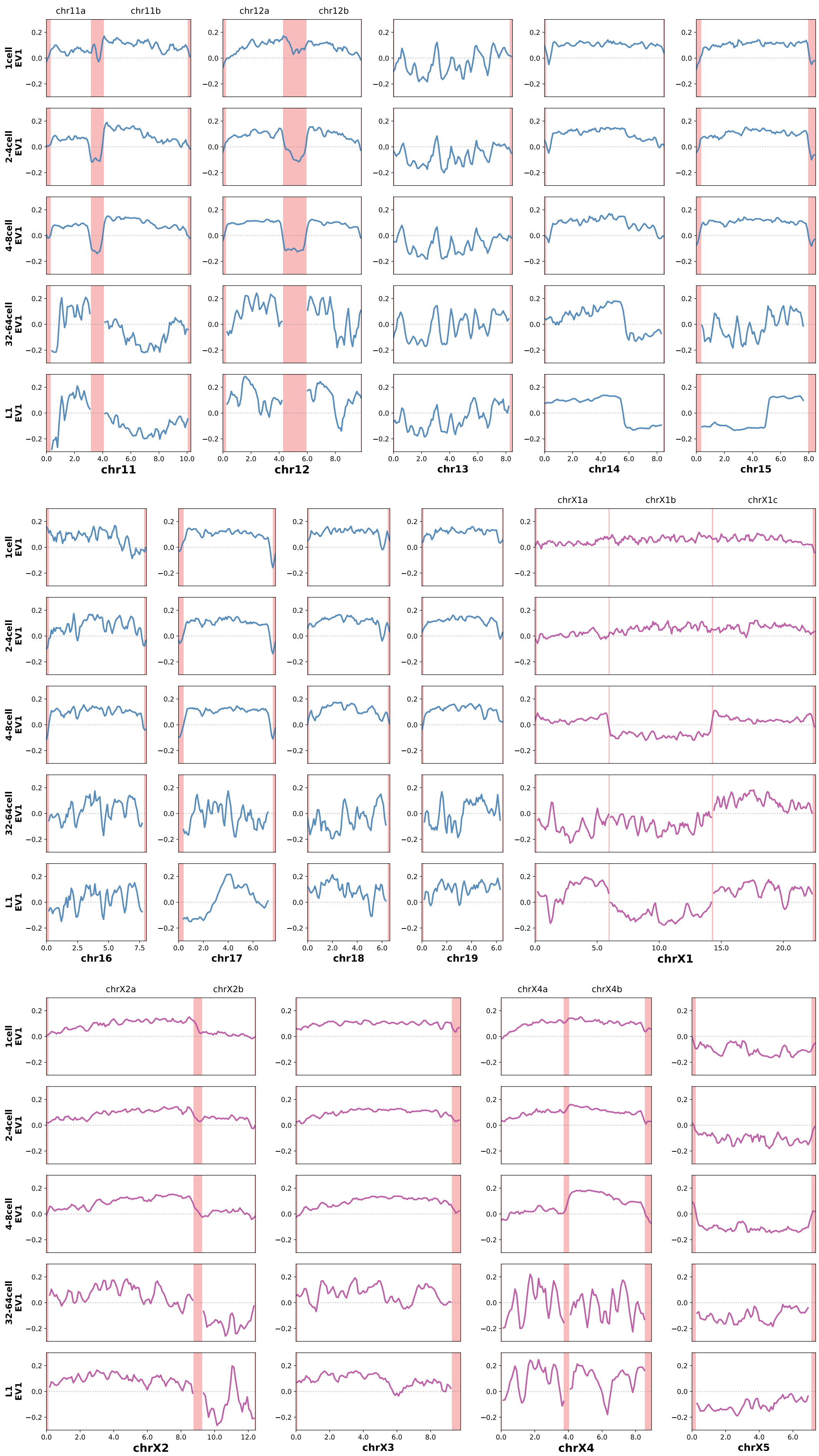


### Figure S7 cont.

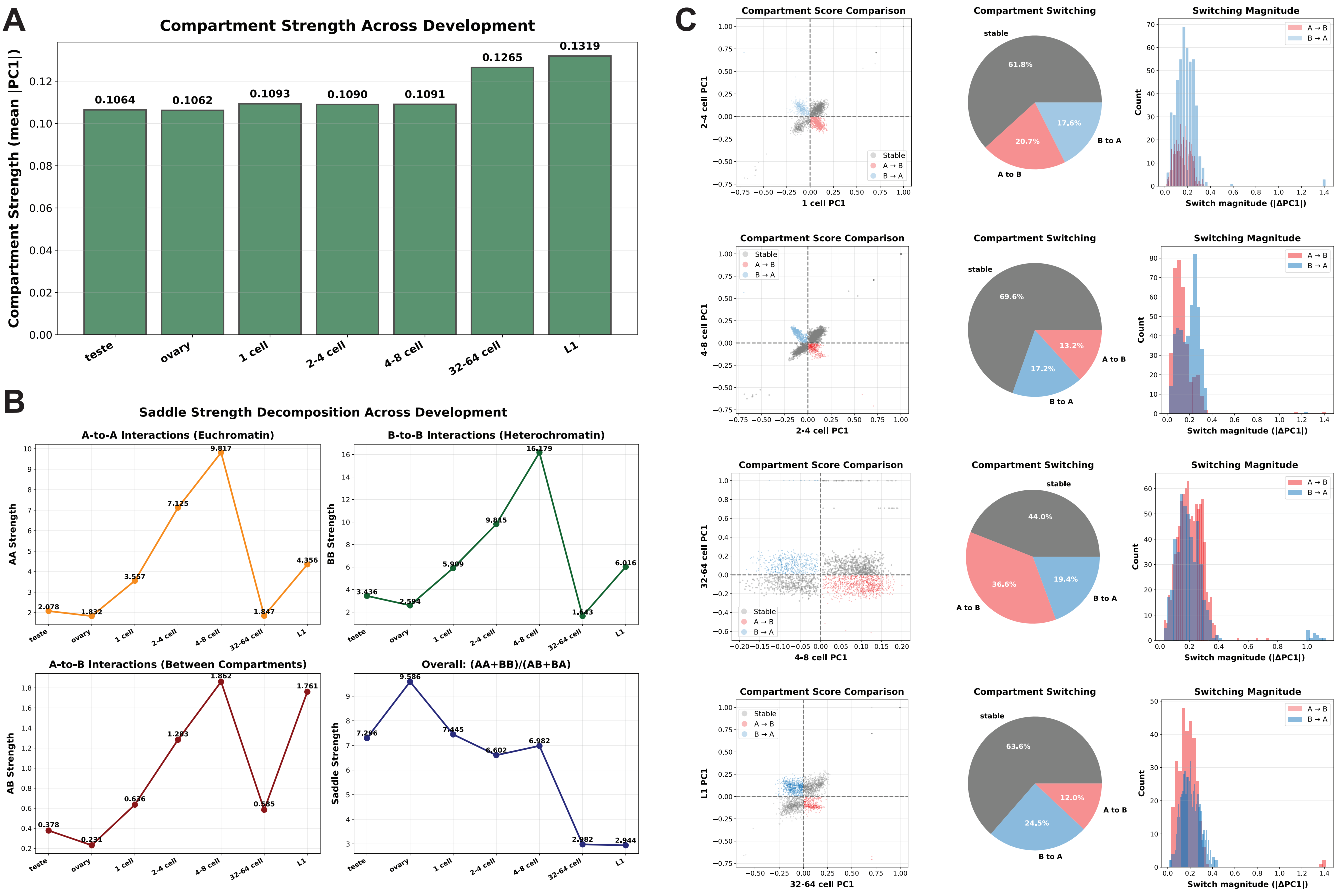

A
Compartment Strength Across Development
Compartment Strength (mean |PC1|)
0.1064
0.1062
0.1093
0.1090
0.1091
0.1265
0.1319
teste
ovary
1 cell
2-4 cell
4-8 cell
32-64 cell
L1
B
Saddle Strength Decomposition Across Development
A-to-A Interactions (Euchromatin)
AA Strength
2.078
1.832
3.557
7.125
9.817
1.847
4.356
B-to-B Interactions (Heterochromatin)
BB Strength
3.436
2.594
5.909
9.815
16.179
1.643
6.016
A-to-B Interactions (Between Compartments)
AB Strength
0.378
0.231
0.636
1.293
1.862
0.585
1.761
Overall: (AA+BB)/(AB+BA)
Saddle Strength
7.296
9.586
7.445
6.602
6.982
2.982
2.944
C
Compartment Score Comparison
2-4 cell PC1
1 cell PC1
Stable
A → B
B → A
Compartment Switching
stable
61.8%
20.7%
17.6%
A to B
B to A
Switching Magnitude
Count
Switch magnitude (|ΔPC1|)
Compartment Score Comparison
4-8 cell PC1
2-4 cell PC1
Compartment Switching
stable
69.6%
13.2%
17.2%
A to B
B to A
Switching Magnitude
Compartment Score Comparison
32-64 cell PC1
4-8 cell PC1
Compartment Switching
stable
44.0%
36.6%
19.4%
A to B
B to A
Switching Magnitude
Compartment Score Comparison
L1 PC1
32-64 cell PC1
Compartment Switching
stable
63.6%
12.0%
24.5%
A to B
B to A
Switching Magnitude


### Figure S8

### *Parascaris* 1 cell (10 hour)

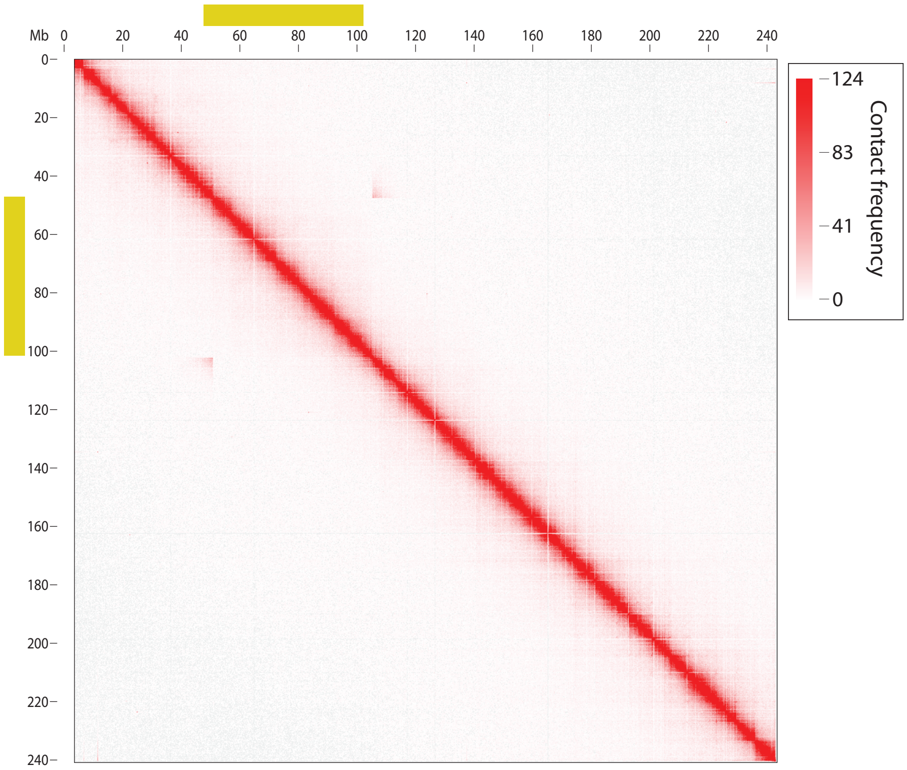


**Figure S9**
